# Hydrogen-Induced Calcium Influx via the TRPC4-TRPC4AP Axis

**DOI:** 10.1101/2025.03.19.644243

**Authors:** Pengxiang Zhao, Han Li, Zisong Cai, Xujuan Zhang, Xiaohu Wen, Ziyi Liu, Shihao Jiang, Zheng Dang, Xue Jiang, Jiateng Wang, Mengyu Liu, Fei Xie, Xuemei Ma

**Author notes:** **Correspondence: Xuemei Ma,**. These authors share first authorship (Pengxiang Zhao, Han Li, Zisong Cai).

## Abstract

**Background:** Calcium ions (Ca²⁺) serve as universal intracellular messengers regulating diverse physiological processes, while dysregulated Ca²⁺ homeostasis triggers cytotoxicity. Molecular hydrogen (H₂) exhibits protective effects against oxidative stress-related pathologies, but its mechanism of action remains incompletely understood. Transient receptor potential canonical 4 (TRPC4) channels and their associated protein TRPC4AP are critical mediators of Ca²⁺ influx ( [Ca²⁺]i), yet their role in H₂-mediated calcium signaling is unexplored. This study investigates the molecular mechanism by which H₂ modulates Ca²⁺ dynamics through the TRPC4-TRPC4AP axis, aiming to establish its therapeutic potential for calcium-related disorders.

**Methods:** The study employed heterogeneous cellular models (e.g., mesenchymal stem cells, neurons, fibroblasts) and in vivo two-photon calcium imaging in C57BL/6J mice. Techniques included CRISPR-Cas9 knockout, siRNA-mediated gene silencing, molecular docking (AlphaFold 3), and protein-protein interaction analysis. Calcium flux was quantified via fluorescence imaging, while mitochondrial integrity and cytoskeletal dynamics were assessed using JC-1 staining, ATPase activity assays, and live-cell imaging. Structural validation of TRPC4-TRPC4AP binding sites utilized mutagenesis and complementation experiments.

**Results:** H₂ selectively enhanced extracellular Ca²⁺ influx via TRPC4-TRPC4AP, with no cytotoxicity or mitochondrial dysfunction observed. Key arginine residues (730Arg-731Arg) in the TRPC4 CIRB domain formed hydrogen-bond networks essential for channel activation. In vivo, H₂ increased neuronal Ca²⁺ transient frequency and amplitude in the primary motor cortex. TRPC4AP knockout abolished H₂-induced Ca²⁺ influx, while mutagenesis of 730Arg/731Arg disrupted channel activity. H₂ also promoted cytoskeletal remodeling and cell motility, dependent on TRPC4AP-mediated Ca²⁺ signaling.

**Conclusions:** This study identifies H₂ as a novel calcium agonist that activates the TRPC4-TRPC4AP axis to regulate extracellular Ca²⁺ influx. The 730Arg-731Arg motif in TRPC4 serves as a critical H₂-sensitive site, enabling dynamic calcium homeostasis without overload. These findings provide a mechanistic basis for H₂-based therapies targeting calcium dysregulation in neurodegenerative, inflammatory, and metabolic diseases, while highlighting TRPC4AP as a pivotal molecular switch for gasotransmitter signaling.

## Introduction

Calcium ions (Ca²⁺) function as a ubiquitous second messenger, mediating various physiological processes through dynamic concentration gradients and complex regulatory mechanisms[1]. Cytoplasmic calcium elevation serves as a universal signaling pathway[2], transmitting signals from the cell surface to the interior, regulating key cellular functions, including gene expression[3], cell cycle progression[4], motility[5], autophagy[6], and apoptosis[7]. In the nervous system, Ca²⁺ is essential for neurotransmitter release, facilitating neuronal communication[8], while in muscle cells, it is critical for contraction by interacting with regulatory proteins like troponin to enable actin-myosin interaction and force generation[9]. Ca²⁺ also activate numerous enzymes, such as thrombin, which is involved in blood clotting[10]. Additionally, extracellular Ca²⁺ maintains the resting membrane potential in excitable cells, such as neurons and muscle cells, and supports proper bone formation[11]. Cytoplasmic Ca²⁺ signals arise from extracellular calcium influx through calcium channels. The endoplasmic reticulum (ER) and mitochondria (Mito) serve as key intracellular calcium reservoirs[12, 13]. In the ER, calcium ions are vital for proper protein folding, and their release via calcium channels contributes to cytoplasmic calcium signaling[14]. Mitochondria play a crucial role in shaping these signals through their calcium transport mechanisms[15].

Intracellular calcium concentrations exhibit dynamic fluctuations, typically ranging from 20 nM to 2 μM, depending on cellular activity[12]. In resting cells, calcium levels are tightly regulated below 200 nM, which is significantly lower than the extracellular concentration (∼1 mM) and the luminal calcium stores in the SR/ER (100–500 μM)[16]. This steep electrochemical gradient is maintained by two primary calcium extrusion mechanisms: the plasma membrane calcium ATPase (PMCA) and the sodium-calcium exchanger (NCX), which cooperate to expel calcium from the cytoplasm[17]. The extracellular space serves as an abundant reservoir for calcium, facilitating its influx into the cytoplasm through various channels, including store-operated, voltage-gated, ligand-gated, and TRP channels. In non-excitable cells, store-operated calcium entry (SOCE) is the predominant pathway, activated upon depletion of ER calcium stores[18]. Additionally, voltage-gated and ligand-gated channels mediate calcium entry in response to changes in membrane potential or ligand binding, respectively[8].

This article focuses on TRP channels, specifically the TRPC family. TRPC channels are permeable to monovalent and divalent cations, including Na⁺, K⁺, Ca²⁺, and Zn²⁺. Among the four subgroups of TRPC channels, three (TRPC1, TRPC4-TRPC5, and TRPC3-TRPC6-TRPC7) form heteromeric complexes in humans[19, 20]. Each TRPC subunit comprises six transmembrane segments that form the cation-permeable pore.The regulation of TRPC channels is influenced by post-translational modifications (e.g., phosphorylation and glycosylation) and specific domains in the N and C termini of the subunits, such as coiled-coil domains, ankyrin repeat domains, calmodulin-binding sites, and binding sites for the inositol 1,4,5-trisphosphate receptor (IP3R1)[21]. TRPC channels can be activated by G protein-coupled receptors (GPCRs) or oxidative stress and are sensitive to changes in intracellular osmolarity or Ca²⁺ levels[20].

Calcium agonists are compounds that increase Ca²⁺ influx into calcium channels in excitable tissues, which is crucial for functions like muscle contraction, vascular regulation, and hormone release[22]. These agonists may have therapeutic potential for treating degenerative diseases, such as amyotrophic lateral sclerosis[23]. However, there are far fewer identified calcium channel agonists compared to the many calcium channel antagonists.

H_2_ is small, non-polar, and highly permeable, enabling it to cross cellular membranes and enter organelles, facilitating interactions with crucial cellular components. Ohta et al. found that 2% H_2_ can effectively reduce cerebral ischemia-reperfusion injury in rats, suggesting a selective antioxidant mechanism[24]. Despite its protective effects against various oxidative stress-related diseases, the traditional antioxidant hypothesis does not fully explain the diverse biological effects of H₂, leading to further exploration of its mechanisms[25].

Our research has revealed that H₂ acts as a calcium ion agonist through the TRPC4 channel and its associated protein, TRPC4-associated protein (TRPC4AP), promoting the influx of extracellular Ca²⁺ into cells and thereby increasing the concentration of calcium ions in the cytoplasm[26]. This effect is widely applicable across various cell types, triggering biological responses in a multitude of cellular contexts. The increase in cytoplasmic calcium ion concentration induced by hydrogen gas stimulation is reversible and does not lead to calcium overload or reduced cell viability. TRPC4AP does not directly constitute the ion channel but serves as a functional partner protein of TRPC4, influencing TRPC4-dependent calcium signaling pathways by regulating its expression levels, subcellular localization, or channel activity.

To elucidate the molecular mechanism of H_2_-induced extracellular Ca²⁺ influx, this study employed siRNA-mediated gene silencing technology for functional validation. We demonstrated that this process is predominantly mediated by the TRPC4 channel localized on the plasma membrane and its interacting partner TRPC4AP. Utilizing the AlphaFold 3 structural prediction platform combined with protein-protein molecular docking techniques, we systematically resolved the three-dimensional conformation of the TRPC4-TRPC4AP complex[27]. Structural simulations and statistical analyses revealed, for the first time, that TRPC4AP regulates channel activity by specifically binding to the CIRB domain (Calmodulin/IP3 Receptor Binding Domain) of the TRPC4 protein. Molecular dynamics investigations further demonstrated that H_2_ modulates channel activity by targeting the CIRB domain within the TRPC4-TRPC4AP complex. Notably, the arginine residues at positions 730 (Arg730) and 731 (Arg731) of TRPC4 play a pivotal role in H_2_-mediated Ca²⁺ transmembrane transport by forming critical hydrogen-bond networks. These findings provide essential structural insights into the gating mechanism of the TRPC4/TRPC4AP complex and its functional role in gas signaling transduction.

This discovery positions TRPC4AP as a unique molecular target for hydrogen therapy, bridging the gap between its observed biological effects and mechanistic understanding. Our findings not only advance H_2_ medicine research but also provide a framework for developing targeted therapies for degenerative diseases, while informing future investigations into H₂ dosing optimization and safety profiles.

## Materials and methods

### Cellular Immunofluorescence Assay

Cells were cultured on glass coverslips within a 6-well plate to the appropriate confluence. Aspirate the medium, wash with PBS, and fix cells with 4% paraformaldehyde for 15 minutes at room temperature. Permeabilize with 0.1% Triton X-100 for 10 minutes, then block with 3% BSA in PBS for 1 hour. Incubate with the primary antibody overnight at 4°C, followed by three washes with PBS. Apply the fluorophore-conjugated secondary antibody for 1 hour at room temperature in the dark, and wash again. Stain nuclei with DAPI for 5 minutes, then mount the coverslips on slides with mounting medium. Analyze the cells using Confocal Fluorescence Microscopy (Nicon, C2, Japan ), ensuring to protect the samples from light to prevent photobleaching.

### Preparation of Cell Samples for TEM

Harvest and Centrifuge cultured cells to form a pellet, then immediately fix the cells in a solution of 2% glutaraldehyde and 2% paraformaldehyde in PBS for 2 hours at 4°C. Wash the pellet with sodium cacodylate buffer, followed by post-fixation in 1% osmium tetroxide. Dehydrate the sample through an ascending ethanol series and transition with propylene oxide. Infiltrate and embed the sample in resin, then polymerize at 60-70°C. Trim and section the embedded block with an ultramicrotome to obtain ultrathin sections. Stain the sections with uranyl acetate and lead citrate, mount them on copper grids, and examine the cellular ultrastructure using a transmission electron microscope (HITACHI, HT7700, Japan).

### Mitochondrial membrane potential detection JC-1

293T cells were seeded at a density of 5×10^3^cells per well in a 96-well plate, with the experimental group receiving culture medium saturated with hydrogen gas. After 24 hours of cell adhesion, when the cell density reached approximately 90%, the cells were used to observe the mitochondrial membrane potential. The culture plate was removed and the cells were gently washed three times with PBS solution to remove impurities from the culture medium. To each well, 200 μL of JC-1 staining working solution with a final concentration of 2 μM was added.The plate was then placed back into the cell incubator and incubated at 37°C for 20 minutes to allow sufficient staining of the mitochondria with JC-1 dye. After incubation, the supernatant was carefully aspirated, and the cells were washed three times with JC-1 staining buffer to remove any unbound dye.200 μL of cell culture medium was added back to each well to maintain the physiological state of the cells. The cells were observed and the fluorescence staining results were recorded using a laser confocal microscope to assess changes in the mitochondrial membrane potential.

### Cell viability assay

The cell viability of the 293T cell line was assessed using the Cell Counting Kit-8 (CCK8). These cells were seeded at a density of 5×10^3^ cells per well in a 96-well plate, and after growing for 24 hours in standard growth medium, the medium was replaced with medium saturated with hydrogen gas. The cell viability was detected using the CCK8 reagent at 2h, 4h, 8h, and 24h time points. The optical density (OD) was measured at 450 nm using a Perkin Elmer microplate reader.

The blank group contained only the culture medium, the control group contained cells that were not treated with saturated hydrogen gas, and the positive control group contained cells treated with 2μM ionomycin to induce calcium overload.

Cell viability = (Abs of the experimental group-Abs of the blank group)/(Abs of the control group-Abs of the blank group) ×100 %.

### Cell death rate assessment and live cell imaging

NucGreen Dead 488 Ready Probes Reagent (ThermoFisher S7020) is a membrane-impermeable stain that selectively labels cells with compromised membranes by binding to DNA and emitting a vivid green fluorescence, without penetrating or affecting the integrity of living cells. This reagent is used to quantify the relative rate of cell death according to the guidelines provided by Thermo Fisher Scientific.

Live cell imaging is a powerful tool for capturing the dynamics of cellular processes and events. Images of 293T cells were captured using the Cytation 5 Cell Imaging Multimode Reader (Biotek Instruments, Inc.,Winooski, VT, USA).Cells were treated with NucGreen Dead 488 Ready Probes Reagent to observe the number of live and dead cells after hydrogen treatment at 2h, 4h, 8h, and 24h time points. The cell imaging data were processed and cell counts were analyzed using Gen5™ Data Analysis Software from Bad Friedrichshalle, Germany.

The control group contained cells that were not treated with saturated hydrogen gas, and the positive control group contained cells treated with 75% ethanol.

### Ca++Mg++-ATPase Activity Assay

Ca++Mg++-ATPase enzymes are widely distributed in plants, animals, microorganisms, and cells, and can catalyze the hydrolysis of ATP to produce ADP and inorganic phosphate. Ca++Mg++-ATPase Assay Kit (Bioss AK266) utilizes the principle of Ca++Mg++-ATPase decomposing ATP to generate ADP and inorganic phosphate, and determines the activity of ATPase by measuring the amount of inorganic phosphate. 293T cells were seeded at a density of 10^5^ cells/mL in T25 flasks and grown in standard growth medium for 24 hours before being replaced with medium saturated with hydrogen gas. After 5 and 10 minutes, the activity of Ca++Mg++-ATPase was determined using the assay kit. The control group contained cells that were not treated with saturated hydrogen gas. The optical density (OD) was measured at 660 nm using a Perkin Elmer microplate reader.

Ca++Mg++-ATPase activity (μmol/h/10^4^) = 0.015 × (A of the test tube - A of the control tube) ÷ (A of the standard tube - A of the blank tube).

### Cell scratch assay

A confluent monolayer of cells was seeded in a 6-well plate. Once confluent, remove the medium and gently scratch the cell layer uniformly across the well with a pipette tip to create a ‘wound.’ Wash the cells with PBS to remove any debris and floating cells. Add fresh culture medium, and return the plate to the Live cell imaging system (Biotek Instruments, Inc., Winooski, VT, USA). Capture images of the wound area at the start and at regular time intervals automaticly. After the experiment, analyze the images to measure the rate of cell migration into the wound area over time. The wound healing rate can be quantified by comparing the wound area at different time points, indicating cell migration and proliferation capabilities.

### Ca^2+^, K^+^, Na^+^ ion probe staining and imaging in live cells

Cells were seeded on glass coverslips in a culture dish to reach the desired confluence. Wash the cells gently with warm Hank’s Balanced Salt Solution (ThermoFisher 14175095) to remove culture medium. Incubate the cells with an ion sensitive fluorescent Probe for Ca^2+^-Fluo-4-AM (5 μM ) diluted in Serum-Free Medium (KeyGEN,KGAF024) ; Probe for K^+^-EPG-4-AM (5 μ M) diluted in Serum-Free Medium(Maokang MX4521) ; Probe for Na^+^-ENG-2-AM (5 μ M ) diluted in Serum-Free Medium(Maokang MX4514); probe for Ca^2+^-Rhod-2-AM (5 μM ) diluted in HBSS(YEASEN 40776ES72) for 10-30 min at 37°C to allow probe uptake. After incubation, wash off excess probe with HBSS or Serum-Free Medium and place the cells in a recording chamber for microscopy. Confocal microscope equipped with a CCD camera (C2, Nicon) was used to capture the fluorescence signal, which indicates ion concentration. Maintain the cells at 37°C and control the atmosphere (CO_2_ level) throughout the imaging process to keep the cells viable, and monitor calcium dynamics in response to stimuli. Analyze the fluorescence intensity changes to assess ion activity within the cells.

### Western blot

Cells were washed, harvested, and centrifuged, and proteins were extracted using a Cell Protein Extraction Kit (San Gong, C006225). All procedures were carried out on ice. After cell lysis, protein quantification was performed using the BCA Protein Assay Kit (Beyotime P0010). Equal amounts of cell protein samples were separated by 10% sodium dodecyl sulfate-polyacrylamide gel electrophoresis (SDS-PAGE) and transferred onto a nitrocellulose membrane (Millipore). The membrane was blocked with a 3% BSA blocking solution for 2 hours at room temperature, then washed three times with TBST, and incubated with the target primary antibody overnight at 4°C. After incubation with a fluorophore-conjugated secondary antibody for 1 hour in the dark, the bands were visualized and captured using the LI-COR® Odyssey Infrared Imaging System.

### Lipofectamine 2000 transfection of siRNA

Cells were seeded at a density of 5×10^3^ cells/well in a 96-well plate and cultured in standard growth medium for 24 hours. On the day of transfection, cells were seeded in an appropriate amount of growth medium without antibiotics to achieve a confluence of 30-50% at the time of transfection. siRNA was diluted in serum-free Opti-MEM (Gibco, 2898884), mixed gently, and Lipofectamine 2000 (ThermoFisher, 11668019) was diluted in Opti-MEM. The diluted siRNA and diluted Lipofectamine 2000 were mixed and incubated at room temperature for 20 minutes to form siRNA-Lipofectamine 2000 complexes. The siRNA-Lipofectamine 2000 mixture was added to each well containing cells and medium. The plate was gently rocked back and forth to mix. Cells were incubated in a 37°C CO_2_ incubator for 48-96 hours, and the medium could be changed after 6-8 hours.

### Molecular Docking

The initial structure of TRPC4 was obtained from the PDB database, with the entrance ID of 7B0J. In the absence of experimentally resolved TRPC4AP structures, we employed AlphaFold we employed AlphaFold to predict the full-length of this associated protein. we utilized SWISSMODEL to refine and complete the TRPC4 protein chain, thus ensuring its functional integrity. In subsequent research, the binding modes and detailed interactions between TRPC4 and TRPC4AP was investigated using protein-protein docking technology.

we generated 2,000 potential docking configurations between TRPC4 and TRPC4AP. Utilizing the PSC (Protein Scoring Function) scoring system, the software performed an automated clustering and scoring of these configurations, ultimately identifying the top 60 TRPC4 and TRPC4 AP binding modes. These initial docking models were based on a rigid docking approach, in an effort to refine these docking configurations ,we employed energy optimization to eliminate potential geometric conflicts in these docking conformations, laying a robust foundation for further experimental validation and clinical applications.

By utilizing advanced protein-docking software, we were not only able to simulate the binding process between TRPC4 and TRPC4AP but could also examine the specific interactions at their interfaces. In a practical application of AlphaFold 3, we input the amino acid sequences of TRPC4 and TRPC4AP into the AlphaFold 3 platform to obtain a model of the TRPC4-TRPC4 AP complex. We analyzed the interfacial interactions within this complex. Notably, an interaction between arginine at position 730 on TRPC4 and a glutamic acid on TRPC4 AP, which received the highest score using the PSC. We have obtained results consistent with our previous protein-protein docking, which confirms that our analysis is correct.

### Construction and Identification of pX459-sgRNA Knockout Plasmids Targeting TRPC4 and TRPC4AP

Based on the genomic information of TRPC4 and TRPC4AP genes from NCBI, the exons were annotated. The nucleotide sequences of the first and second exons of the target genes were input into the sgRNA online design website http://crispr.mit.edu. Two sgRNA sequences with the lowest off-target rates were selected. After synthesis by Suzhou Hongxun Biotechnology Co., Ltd., they were named: TRPC4 sgRNA-1 and TRPC4 sgRNA-2; TRPC4 AP sgRNA-1 and TRPC4 AP sgRNA-2. The sgRNA primers were annealed to form double-stranded DNA through a programmable annealing process. The vector pX459 was linearized after digestion with BbsI enzyme and then recovered. T4 DNA ligase was used to ligate the annealed sgRNA double-stranded DNA with the linearized pX459 at room temperature to obtain the recombinant plasmids, which were then sequenced and identified for correctness before being used in subsequent experiments.

### Preparation and Selection of 293T Cells with TRPC4 and TRPC4AP Gene Knockout

Seed HEK293T cells into a 6-well cell plate, and when the cell confluence reaches 80%, transfect the 293T cells with the pX459 plasmid carrying TRPC4 and TRPC4AP sgRNA. Meanwhile, transfect with empty plasmid pX459 in the same manner as a negative control. After 36 hours, perform pressure screening with 1μg/mL puromycin. After two rounds of puro selection, once the control group cells have all died, digest the experimental group cells into single cells, and use the limited dilution method to sort out monoclonal cells and expand the culture. After passaging the cell line, freeze and store for future use.

### Identification of Gene and Protein Knockout in 293T Cell Lines with TRPC4 and TRPC4AP Gene Knockout

Collect monoclonal cell strains and number them. Extract the genomic DNA from the cells according to the instructions of the DNA extraction kit. Based on the genomic sequences of TRPC4 and TRPC4AP targeted by sgRNA, design specific identification primers according to the intron sequences on both sides of the corresponding exons. Use these primers and the extracted cellular genomic DNA as templates to perform PCR amplification. The amplified products are sequenced and aligned to detect the knockout effect of the target genes in the obtained monoclonal cell strains.

### Animal Studies

Male C57BL/6J mice (8 weeks old) were obtained from Charles River Laboratories (Beijing). Mice were maintained in a 12-hour light/dark cycle at 25°C with free access to food and water. All animal experiments were approved by the Institutional Animal Care and Use Committee (IACUC) and were conducted in accordance with the guidelines for the ethical treatment of animals. To monitor Ca²⁺ influx in the primary motor cortex (M1), C57BL/6J mice were stereotactically injected with adeno-associated virus (AAV) expressing Syn1-driven jGCaMP7 (AAV-Syn1-jGCaMP7).

Mice were anesthetized using isoflurane 2% in oxygen and positioned in a stereotactic frame. The scalp was shaved and disinfected with ethanol. A midline incision was made, and a small craniotomy (1-2 mm in diameter) was performed at the coordinates (X=2 mm lateral, Y=0.8 mm anterior, Z=2.4 mm depth) relative to bregma, to target the primary motor cortex (M1). AAV-Syn1-jGCaMP7 (titer: 1 × 10^13^ vg/mL) was injected at a rate of 0.2 µL/min using a microinjection pump (UltraMicroPump3 (UMP3)). A total of 50 µL of AAV vector was delivered into M1. The needle was left in place for an additional 5 minutes to allow for viral diffusion, and the incision was closed with sutures. Mice were allowed to recover for 2-3 weeks to ensure proper viral expression and cellular uptake.

### Two-Photon Microscopy for In Vivo Calcium Imaging in the Primary Motor Cortex (M1)

Experimental Setup and Preparation To monitor real-time calcium dynamics in the primary motor cortex (M1) of C57BL/6J mice infected with AAV-Syn1-jGCaMP7, in vivo two-photon imaging was performed as follows:

Animal Preparation . After the recovery period post-surgery and sufficient expression of jGCaMP7 in the M1 region, mice were anesthetized with isoflurane (2% for induction, 1-1.5% for maintenance) and placed onto a custom imaging platform. Core body temperature was maintained at 37°C using a heating pad.

Cranial Window and Imaging Setup. A cranial window above the primary motor cortex was prepared by carefully removing the overlying skin and gently clearing the bone over M1. A round coverslip was placed on the exposed cortical surface to protect the brain tissue and reduce motion artifacts. The window was sealed with dental cement (C&B Metabond) to secure the coverslip in place.

Imaging Preparation. The mouse was placed under the objective lens of the two-photon microscope (Olympus FV1000, Leica SP8), and the brain region of interest was positioned in the imaging field. The system was equipped with a Ti:sapphire laser for two-photon excitation (900 nm), and the emitted fluorescence from jGCaMP7 was collected through a 510–550 nm bandpass filter.

### Bulk-RNA seq and analysis

The raw RNA-seq data were processed using a standard pipeline by Annoryoda, and were then deeply analyzed and visualized using R (version 4.4.1). The script utilized in this study is provided in the supplemental materials. Differential expression genes were screened based on the criteria of an adjusted P-value < 0.05 and an absolute fold change > 2. Principal component analysis (PCA) was conducted to assess the overall variance in the dataset, and hierarchical clustering was performed to group samples according to their gene expression patterns. Heatmaps of the differentially expressed genes were generated using the pheatmap package. Gene ontology (GO) and Kyoto Encyclopedia of Genes and Genomes (KEGG) enrichment analyses were carried out on the differentially expressed genes using the clusterProfiler package, aiming to identify the biological processes and pathways that are enriched in the data. GSEA analysis was also performed to demonstrate the expression trends of all genes.

In detail, we conducted individual K-Means fuzzy clustering analyses to delineate the transcriptomic profiles of DEGs in MSCs exposed to H_2_ for 2 hours (MSCs-2h-H_2_) versus control conditions (MSCs-2h-Ctrl), and after 24 hours (MSCs-24h-H_2_) versus control conditions (MSCs-24h-Ctrl). Subsequently, we performed an integrated K-Means clustering on the top 5000 DEGs identified from the aforementioned pairwise comparisons to elucidate the overarching transcriptional dynamics. GO enrichment analysis was subsequently applied to each distinct cluster to infer functional annotations and biological processes that were significantly enriched. Furthermore, we executed a temporal transcriptome analysis to track the evolution of gene expression patterns over the specified time intervals, providing insights into the temporal regulation of MSCs in response to H_2_.

## Results

### H_2_ promotes the accumulation of Ca^2+^ in the cytoplasm

Ca²⁺, functioning as master intracellular secondary messengers, orchestrate cellular signaling networks and physiological processes through spatiotemporal concentration gradients. This study unveils a novel molecular mechanism whereby H₂ specifically activates cytosolic Ca²⁺ signaling in a dynamically reversible manner. Systematic interrogation across eight heterogeneous cellular models (Fig 1, Fig 2). Including human umbilical cord-derived mesenchymal stem cells (MSCs) Fig. 1A; bone marrow-derived mesenchymal stem cells (iBMSCs) Fig. 2A, mouse osteoblasts (MC3T3-E1) Fig. 2B; mouse myoblasts (C2C12) Fig. 2C,; mouse fibroblasts (NIH-3T3) Fig. 2D,; human skin fibroblasts (ESF) Fig. 2E, and rat neuronal cells (PC12) Fig. 2F.

**Fig. 1.**
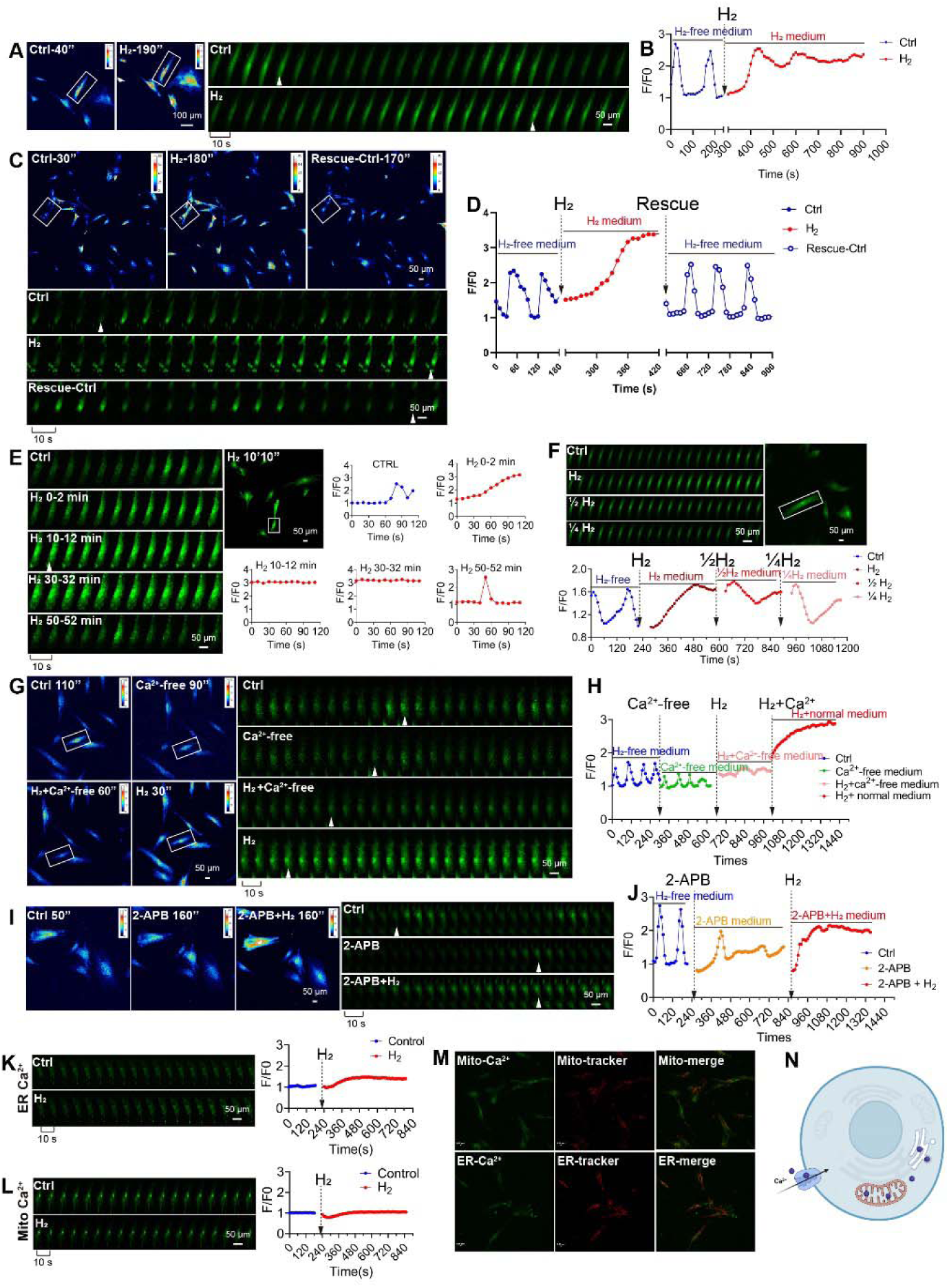
Dynamic time-lapse Ca^2+^ imaging by the Stimulation of molecular H_2_ induces [Ca^2+^]i oscillations in MSCs. A. Pseudocolor and time series images of the [Ca^2+^]i changes under H_2_-free and H_2_-medium conditions. B. Fluo 4 averaged F/F0 trace in imaging H_2_-free (blue) and H_2_-medium conditions (red). C & D, Pseudocolor (C left, upper) and time series (C left, lower) images and the Fluo-4 averaged ΔF/F0 trace of the [Ca^2+^]i changes under H_2_-free, H_2_-medium, and then back to H_2_-free conditions. E. Different time period of Fluo-4 images and averaged F/F0 traces in H_2_-induced MSCs [Ca^2+^]i during extended time-lapse imaging. F. Time-lapse Ca^2+^ images and Fluo-4 averaged F/F0 trace of MSCs under H_2_-free, H_2_-medium (Saturated H_2_ medium), 1/2H_2_-medium, and 1/4H_2_ medium. G & H, Pseudocolor (G left) and time series (G right) images and the Fluo4 averaged F/F0 trace of the [Ca^2+^]i changes under H_2_-free, Ca^2+^-medium, H_2_+Ca^2+^free-medium, and then back to H_2_-medium conditions. I & J, Pseudocolor (I left) and time series (I right) images and the Fluo4 averaged F/F0 trace of the [Ca^2+^]i changes under H_2_-free, 2-APB-medium, and then back to 2-APB+H_2_-medium. K & L, Time series images and the averaged F/F0 trace of the [Ca^2+^]i changes in ER (K) and Mito (L) under H_2_-free and H_2_-medium. M, Co-localization of Mito-Ca^2+^ and ER-Ca^2+^ with their trackers, respectively. N, simplified model illustrating the promotion of [Ca^2+^]i by H_2_ from extracellular. White arrows in A, C, E, G, and I indicate cells at distinct time points within the time-lapse capture. Sampling rate, 10 sec.

**Fig.2.**
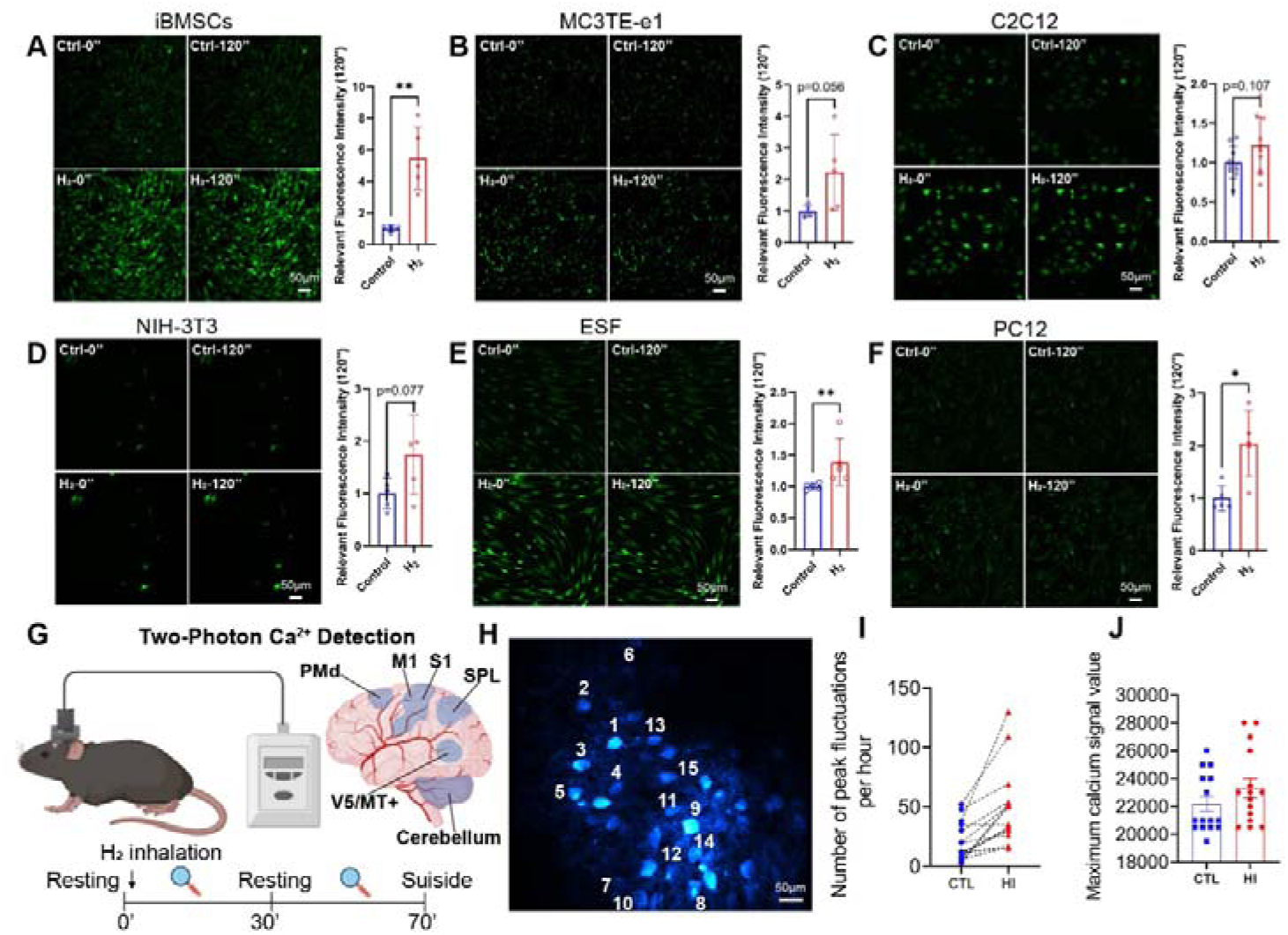
In-vitro and In-vivo assays confirmed H_2_ promoted [Ca^2+^]i in varous cells. A-F, H_2_ stimulates calcium ion influx in various cells, including mouse bone marrow-derived mesenchymal stem cells (iBMSCs), mouse osteoblasts (MC3T3-E1), mouse myoblasts (C2C12), mouse fibroblasts (NIH-3T3), human skin fibroblasts (ESF), and rat neuronal cells (PC12); The left panels show random field fluorescence images of Ca^2+^ at 0 and 120 seconds, respectively, and the right panels display the relative fluorescence intensity statistics at 120 seconds. G, upper left, common window of mouse brain preparations for in vivo two-photon Ca^2+^ imaging; upper right, six distinct functional areas of the mouse cerebral cortex, including the monitored Primary Motor Cortex (M1) region.; lower, schematic diagram of the animal experimental procedure. H, randomly selected 15 neurons in the M1 region. I, comparison between Conctrl group (CTL) and H_2_ inhalation group (HI) in the number of peak fluctuations per hour. J, comparison between CTL and HI groups in the maximum calcium signal value. *p < 0.05, **p< 0.01.

We revealed that H₂-induced Ca²⁺ mobilization exhibits graded concentration-dependent responsiveness. Taking human umbilical cord derived mesenchymal stem cells (MSCs) as an example, a paradigmatic model, H₂ exposure triggered rapid Ca²⁺ elevation (ΔF/F₀ = 2.8 ± 0.3 within 30 s; Fig 1A, Fig 1B), with signal intensity returning to baseline upon H₂ removal (Fig 1C, Fig 1D, Figure S1A). Strikingly, spontaneous resurgence of calcium oscillations—a conserved signaling modality characterized by periodic Ca²⁺ fluctuations was observed approximately 50 minutes post-H₂ withdrawal (Fig. 1E). Concentration-gradient experiments further demonstrated a 42.7% ± 5.1% reduction in Ca²⁺ fluorescence intensity when H₂ concentration decreased from saturation (1.6 ppm) to half-saturation (0.8 ppm) (Fig. 1F). Crucially, this regulatory mechanism operated independently of oxygen tension (Figure. S1B), suggesting H₂ modulates calcium homeostasis through non-canonical pathways, potentially involving TRPC4/TRPC4AP complex targeting.

### Extracellular Ca^2+^ account for H_2_-induced cytoplasmic Ca^2+^ accumulation

Cytosolic Ca²⁺ accumulation may originate from three principal sources: extracellular influx, release from ER calcium reservoirs, or mitochondrial calcium discharge. Our investigations demonstrated that H₂ had no significant effect on cytosolic Ca²⁺ under Ca²⁺-free medium conditions. Remarkably, upon reintroduction of Ca²⁺-containing medium, H₂ regained its capacity to accumulate Ca²⁺ in the cytoplasm, with the fluorescence intensity ratio (F/F₀) recovering to 92.4% ± 6.7% (Fig. 1G; Fig.1H). To investigate the role of ER calcium stores in H₂ induced calcium signaling, we employed pharmacological calcium depletion using 2-aminoethoxydiphenyl borate (2-APB). Experimental results demonstrated that even under ER calcium store-specific depletion conditions, H₂ still significantly elevated cytosolic Ca²⁺ concentrations (Fig. 1I; Fig.1J). Using fluorescence probes specifically targeting ER and mitochondrial calcium, quantitative analysis via dual-channel ratiometric fluorescence imaging revealed no significant changes in calcium levels within the ER or mitochondria under H₂ exposure. Colocalization of mitochondrial calcium (Mito-Ca²⁺) and endoplasmic reticulum calcium (ER-Ca²⁺) with their respective calcium indicators(Fig. 1M). A simplified model illustrating how H₂ promotes the increase of intracellular calcium ([Ca²⁺]i) through the influx of extracellular calcium (Fig. 1N).Statistical analysis of calcium source contributions based on fluorescence intensity further indicated that the observed increase in intracellular calcium signals primarily originated from extracellular calcium influx, while ER calcium release exhibited minimal contribution. These findings confirm the dominant role of extracellular calcium entry in mediating H₂-regulated calcium homeostasis.

### Broad-Spectrum Ca^2+^ Accumulation in the Cytoplasm Induced by H_2_

To validate the broad-spectrum effects of H₂ , we initially investigated its impact on human umbilical vein endothelial cells (HUVECs) (Figure S2). Our findings revealed that hydrogen gas induced cytoplasmic calcium ion (Ca²⁺) accumulation in HUVECs, with a more rapid decline rate compared to mesenchymal stem cells (MSCs) (Figure. S2A, Figure. S2B). This hydrogen-induced calcium elevation exhibited reversibility, as calcium concentrations returned to baseline levels following replacement with hydrogen-free standard medium (Figure. S2A, Figure. S2B). Notably, similar to observations in MSCs, the hydrogen-triggered calcium increase in HUVECs showed minimal dependence on endoplasmic reticulum (ER) calcium stores, with extracellular calcium influx playing a predominant role. This conclusion was supported by both ER calcium store inhibitor experiments (Figure. S2C, Figure. S2D) and calcium-free medium tests (Figure. S2E, Figure. S2F) in HUVECs.

Expanding our investigation to diverse cell types, we observed varying magnitudes of hydrogen-induced cytoplasmic calcium elevation across multiple cellular models, including iBMSCs (Fig. 2A); MC3TE-e1 (Fig. 2B); C2C12 (Fig. 2C), NIH-3T3 (Fig. 2D), ESF (Fig. 2E), and PC12 (Fig. 2F). These collective results demonstrate the universal efficacy of hydrogen gas in promoting calcium influx across distinct cell types.

To further validate the regulatory effect of hydrogen on intracellular Ca²⁺ in vivo, we employed two-photon in vivo calcium imaging (Fig. 2G). C57BL/6J mice received stereotaxic injections of AAV-Syn1-jGCaMP7 into the primary motor cortex (M1) for neuron-specific calcium indicator expression. Quantitative analysis of 15 neuronal cells in the M1 region (Fig. 2H) revealed that hydrogen inhalation (HI) significantly increased the frequency of calcium transients compared to the control group (CTL), with mean hourly peak counts of 50 versus 25 respectively (Fig. 2I). Moreover, hydrogen exposure enhanced the maximum calcium signal amplitude (CTL: 22,000 ± 500 vs. HI: 23,000 ± 450) (Fig. 2J). These in vivo findings closely aligned with our previous cellular experimental observations, collectively demonstrating that hydrogen effectively promotes calcium influx in both in vivo and in vitro systems.

### H_2_ Promotes Ca^2+^ Influx without Inducing Mt Ca^2+^ Overload

High levels of intracellular Ca^2+^ can lead to severe cytotoxicity, particularly in the regulation of mitochondria, where Ca^2+^ enter and exit through a variety of specialized channels and transporters in a tissue-specific manner. Relatively low levels of Ca^2+^ concentration are necessary for mitochondrial energy metabolism, while excessive mitochondrial calcium can directly destroy energy metabolism and lead to apoptosis. Calcium overload is an intermediate state that can disrupt oxidative phosphorylation and is associated with multiple organ dysfunctions. We used ionomycin as a positive control for calcium overload and performed CCK8 assays, which showed that cell viability of MSCs at 24-hour, and 293T at 4-hour, 8-hour, and 24-hour time points was higher in the H_2_ group than in the ionomycin group, with no significant difference from the control group (Fig. 3A, Fig. 3G).

**Fig. 3.**
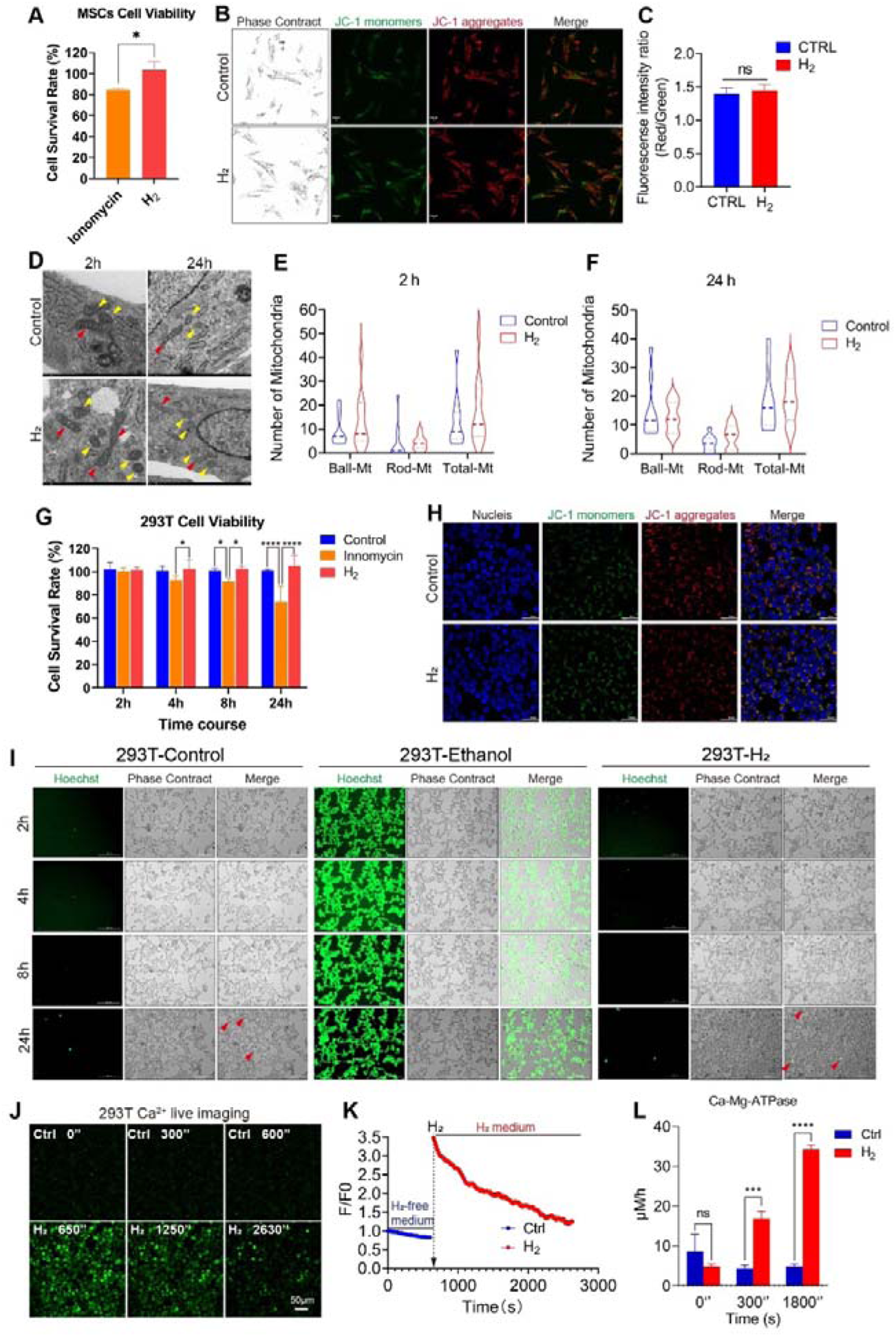
H_2_ promotes [Ca^2+^]i without altering cellular activity or Ca^2+^ overload. A & G, The cell viability of MSCs (A) and 293T (G) remained unaffected by H_2_ exposure, in contrast to the significant cytotoxicity induced by the Ca^2+^ ionophore lonomycin. Mitochondrial membrane potential (MMP) measurement by JC-1 dye in B, C (MSCs) and H (293T). D, Mitochondrial morphological changes in MSCs induced by H_2_ at 2 and 24 hours post-treatment according to the transmission electron microscopy (TEM) detection. E & F, Comparison of ball, rod-shaped, and total mitochondrial counts in MSCs after 2 and 24 hours of H_2_ treatment. I, Live cell imaging of 293T cell death staining following 24 hours of H_2_ treatment. J & K, Time-lapse live cell imaging of H_2-_promoted [Ca^2+^]i dynamics and F/F0 fluorescence intensity analysis. L, Temporal analysis of Ca-Mg-ATPase activity in 293T cells following H_2_ treatment. *p < 0.05, ***p< 0.001, ****p< 0.0001, ns: no significance.

JC-1 staining indicated that after 24 hours of H_2_ treatment, the mitochondrial membrane potential was not significantly different from the control group (Fig. 3B, Fig.3C, and Fig.3H), indicating that mitochondrial function was unaffected. After H_2_ treatment for 2 hours and 24 hours, there was a noticeable increase in the number of rod-shaped mitochondria and the total mitochondrial count in MSCs (Fig.3D, Fig.3E and Fig.3F), suggesting an enhancement of mitochondrial energy metabolism by H_2_. H_2_ treatment, which is a single treatment, can affect Ca^2+^ influx as the H_2_ concentration in the culture medium decreases. After a single treatment of 293T cells with H_2_ for 2630 seconds, the intracellular Ca^2+^ concentration basically returned to the level of the control group (Fig. 3J, Fig. 3K), suggesting that Ca^2+^ may also be actively pumped out. Further measurement of Ca-Mg-ATPase activity at different time points revealed that the enzyme activity in the H_2_ group increased over time, showing a gradually rising trend (Fig. 3L), and reached more than six times that of the control group after 30 minutes. Another channel for calcium extrusion at the cell membrane is NXC, and since the Na^+^ concentration did not change significantly (Figure. S3B, Figure. S3D), the role of this channel in calcium extrusion is excluded. In addition, after a single treatment with H_2_, the endoplasmic reticulum Ca^2+^ slightly increased (Fig. 1 K), possibly because the ER stored the excess Ca^2+^ from the cytoplasm. In summary, the calcium influx induced by hydrogen treatment can be avoided by extrusion through cell membrane-related channels or storage in the ER, preventing excessive calcium in the cytoplasm and thus avoiding calcium overload.

### TRPC4 Channel: A Pathway for H_2_-Promoted Ca^2+^ Influx and Its High Dependency on TRPC4AP Protein

Through transcriptome sequencing analysis, we systematically compared FPKM values of calcium channel-related genes in MSC membranes, revealing predominant expression of T-type, L-type, and N-type voltage-gated calcium channels (VGCCs) (Figure. S4A). Among transient receptor potential (TRP) channels, TRPM4, TRPM7, TRPV2, TRPC1, and TRPC4 showed prominent expression (Figure. S4B), with TRPC4AP being the major expressed TRPC4-associated protein (Figure. S4B). Regarding store-operated calcium entry (SOCE) components, Orai1, Orai2, and STIM1 were identified as the principal expressed proteins (Figure.S4C).

To identify hydrogen-dependent calcium channels, we conducted inhibitory and siRNA interference experiments targeting these identified channels. Pharmacological inhibition of T-type, L-type, or N-type VGCCs demonstrated minimal impact on H₂-enhanced calcium influx (Figure. S4D, Figure. S4E and Figure. S4F). Furthermore, H₂-mediated effects remained unaffected by TRPM4/7, TRPV2, Orai1/2, or STIM1 inhibition (Figure.S4 H, Figure.S4J, Figure.S4K, Figure.S4L, Figure.S4N, Figure.S4O, Figure.S4P, Figure.S4Q). Notably, both pan-TRPC family inhibitors (Figure.S4G) and specific TRPC4/5 antagonists (Figure.S4I) effectively abolished hydrogen-induced calcium influx.

The TRPC1, TRPC4, and TRPC5 channel proteins frequently assemble into heterotetrameric complexes, with TRPC4 exhibiting high homology to TRPC5. Gene silencing experiments demonstrated that H_2_-mediated Ca^2+^ influx does not require TRPC5 or TRPC1 (Figure.S4I, Figure.S4L), but TRPC4 silencing markedly reduced H_2_-induced effects (Fig. 4A, Fig. 4B). Importantly, H_2_-triggered Ca^2+^ influx remained unaffected by Giα knockdown, confirming its mechanism operates independently of Giα signaling (Figure. S4P).

**Fig. 4.**
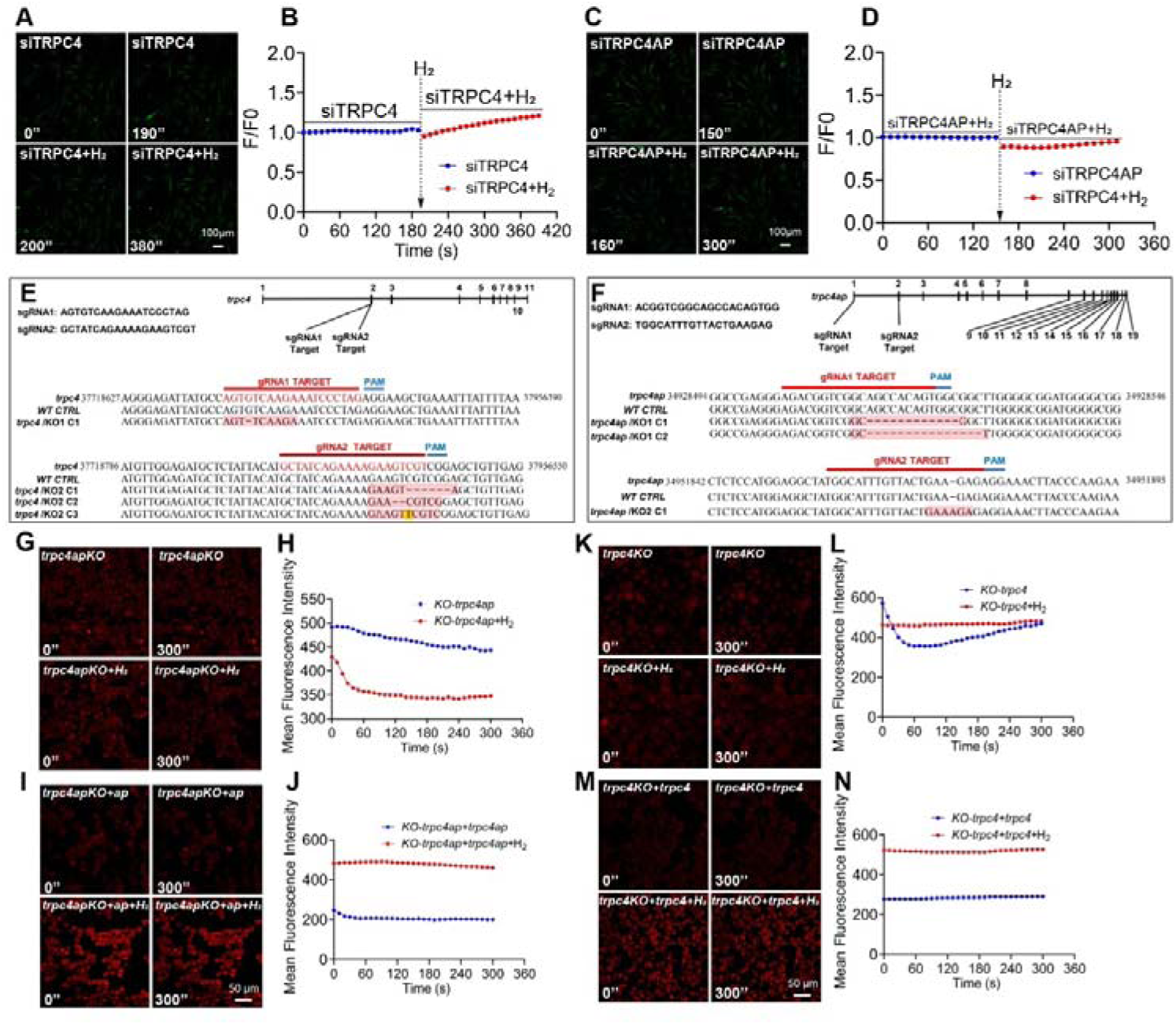
The [Ca^2+^]i induced by H_2_ is abolished following the knockdown or knockout of *trpc4* or *trpc4ap* genes. A & C siRNA-mediated TRPC4 and TRPC4ap knockdown abrogated the [Ca^2+^]i triggered by H_2_. E & F, Confirmation of trpc4 and trpc4ap knockout in monoclonal 293T cell lines generated by CRISPR-Cas 9. For each sgRNA, the target sequence is shown. The mutant site is highlighted by red color, and the protospacer adjacent motif (PAM) is underlined. Sequences of knockout cells were determined by next-generation sequencing. G & K, [Ca^2+^]i induced by H_2_ vanished following the knockout of trpc4 or trpc4 ap. I & M, [Ca^2+^]i induced by H_2_ recovered after trpc4 or trpc4ap overexpression in the trpc4 or trpc4ap knockout cell lines. B, D, H, J, L, & N, graphs showing the Fluo4 averaged F/F0 trace of the [Ca^2+^]i changes under different conditions.

Multichannel inhibition analyses identified TRPC4 and TRPC4AP as principal mediators of H_2_-driven Ca^2+^ influx (Fig. 4). Genetic ablation of TRPC4AP completely eliminated H_2_-mediated regulation of Ca^2+^ influx (Fig. 4C, Fig. 4D), establishing TRPC4AP as a crucial molecular target for H_2_ action. TRPC4AP interacts directly with TRPC4 while maintaining associations with TRPC1 and TRPC5, implying its functional involvement in organizing heteromeric complexes among these proteins.

To delineate mechanistic contributions of TRPC4 and TRPC4AP in H_2_ signaling, CRISPR-Cas9-generated TRPC4 and TRPC4AP knockout 293T cell lines were established (Fig.4E, Fig.4F), with sequencing data confirming knockout efficiency (Figure. S5). Both TRPC4 (293T^trpc4−^) and TRPC4AP(293T^trpc4ap−^) cells exhibited complete loss of H_2_-induced Ca^2+^ responses (Fig.4G, Fig.4H, Fig.4K, Fig.4L). Functional restoration through reintroduction of wild-type TRPC4 (wt-trpc4) or TRPC4AP (wt-trpc4ap) reinstated calcium signaling responsiveness to H_2_ in respective knockout models (Fig. 4I, Fig. 4J, Fig. 4M, Fig. 4N). These findings provide definitive evidence for the essential roles of TRPC4 and its interacting partner TRPC4AP in mediating H_2_-dependent calcium signaling transduction pathways.

### Molecular Docking Prediction and Validation of the TRPC4-TRPC4AP Protein Binding Site

To investigate the interaction between TRPC4 and TRPC4AP, we performed protein-protein docking analyses using the structural file of TRPC4 (PDB ID: 7b0j) and an AlphaFold-predicted model of TRPC4AP. From 2,000 potential docking conformations, we selected the top 60 TRPC4-TRPC4AP interaction patterns based on computational scoring metrics. Structural alignment revealed a unique cytokine-induced SH_2_ domain-binding (CIRB) domain in TRPC4 (Fig. 5A, Fig. 5B), a critical regulatory element in cellular signaling. This domain typically facilitates interactions with SH_2_ domains to coordinate essential cellular processes, including growth regulation, differentiation programming, migration control, and survival mechanisms.

**Fig. 5.**
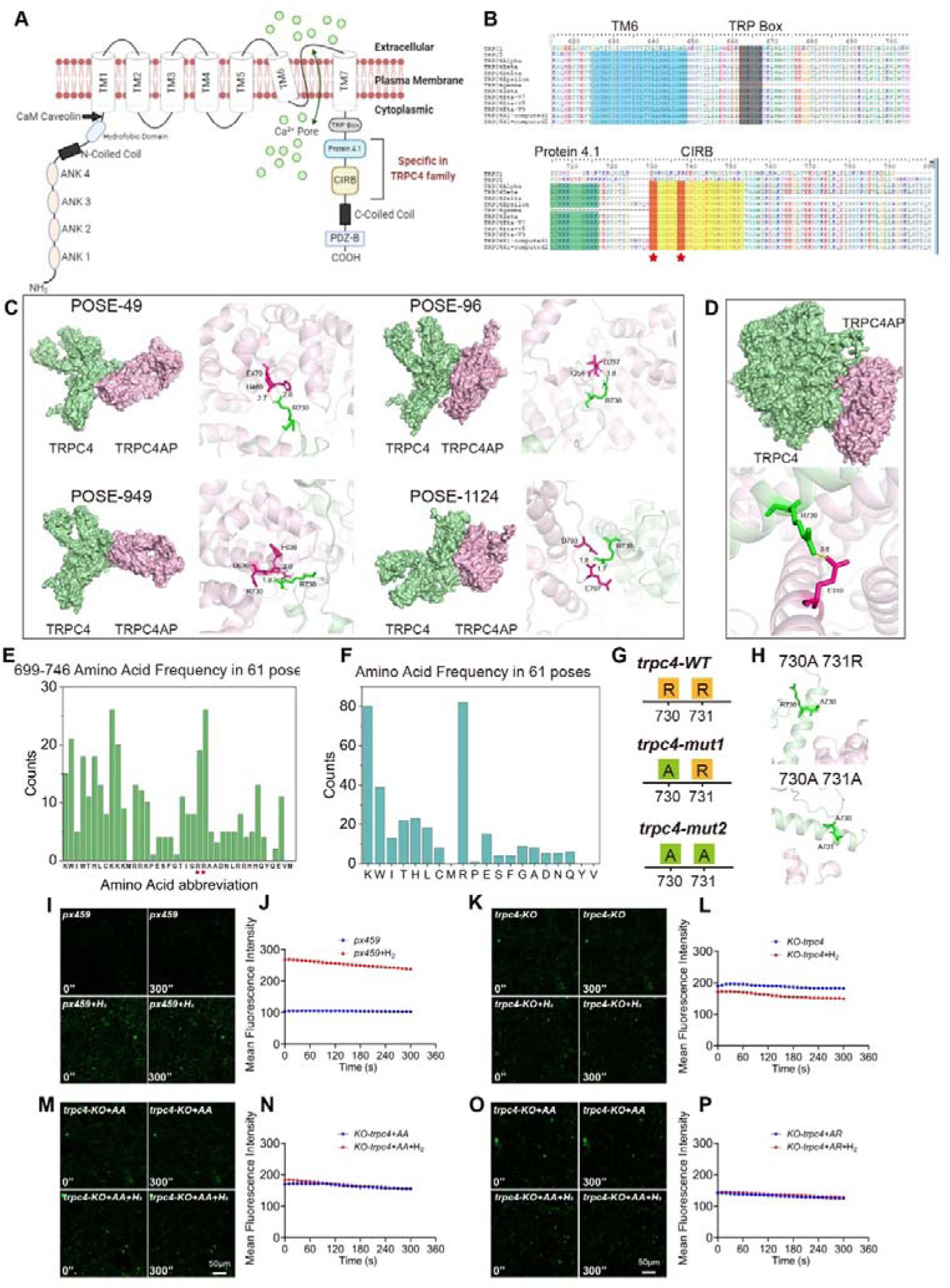
Molecular docking and mutational analyses confirm that TRPC4-TRPC4ap interaction is essential for H_2_-induced [Ca^2+^]i. A, Diagram of the structure of an TRPC4 Ca^2+^ channel including the specific motifs for TRPC4 family. B, Amino acid sequence alignment of TRPC1, TRPC5, and 11 subtypes of the TRPC4 family within the TM6, TRP Box, Protein 4.1, and CIRB domains. The specific arginine residues within the TRPC4 family are highlighted, and the areas of interest (arginine residues) in the CIRB motif are indicated by red pentagrams in the sequence alignment. C & D, Molecular docking utilizing advanced protein-docking software (C) and the AlphaFold 3 platform (D) confirmed the interaction binding sites of TRPC4 and TRPC4ap, with consistent prediction outcomes. E & F, Amino acid frequency analysis at various positions (E) and total count (F) from the top 61 TRPC4-TRPC4ap combination patterns with the highest scores, double Arginine residues are indicated with red pentagram symbol. G, Schematic representation of the wild-type and two mutant forms of the TRPC4 protein at positions 730 and 731 at the C-terminus. H, Molecular docking utilizing the AlphaFold 3 predicted the binding affinity of two TRPC4 mutants with TRPC4ap at positions 730 and 731, indicating a loss of binding interaction. I-P, [Ca^2+^]i induced by H_2_ and the Fluo4 averaged F/F0 trace of the [Ca^2+^]i changes under different conditions, including blank vehicle, trpc4-KO, trpc4-KO+Arg^730^Arg^731^, trpc4-KO+Ala^730^Arg^731^.

Detailed analysis of TRPC4-TRPC4AP complex formation demonstrated that the CIRB domain of TRPC4 mediates interaction with TRPC4AP. Through amino acid frequency mapping of TRPC4 conserved regions (Fig. 5E, Fig. 5F) combined with computational modeling, we identified two adjacent arginine residues (positions 730 and 731) in the C-terminal region of the CIRB domain as primary interaction sites (Fig. 5C). Parallel predictive analyses using AlphaFold 3 and the same docking software revealed consistent binding patterns, with the interaction between TRPC4 arginine-730 and a glutamate residue on TRPC4AP achieving the highest protein scoring function (PSC) value (Fig. 5D). Both methodologies produced compatible spatial configurations and energetically favorable interaction profiles[27].

To elucidate the molecular basis of TRPC4-TRPC4AP interaction, we conducted systematic mutagenesis targeting arginine residues at positions 730 and 731 within the CIRB motif. AlphaFold 3-based structural predictions demonstrated complete disruption of protein-protein interaction upon single (R730A) or double (R730A/R731A) mutations (Fig. 5G, Fig. 5H), establishing the essential role of these conserved residues in complex formation.

To verify the predictions made by molecular docking, we constructed amino acid mutation vectors in the px459 system, guided by human amino acid preferences. We mutated the wild-type arginines (wt730^Arg^731^Arg^) in the CIRB motif to 730^Ala^731^Arg^ and 730^Ala^730^Ala^ to investigate the dependence of H_2_ on the TRPC4-TRPC4AP binding site. The results demonstrated that, in the presence of H_2_, Ca^2+^ concentration in the px459 pressure screening control group increased more than twofold (Fig. 5I, Fig. 5J), while the *trpc4*KO cell line showed no response to H_2_-induced Ca²⁺ influx (Fig. 5K, Fig. 5L). Upon complementing the *trpc4*KO cell line with px459-730^Ala^730^Ala^ (Fig. 5M, Fig. 5N) and px459-730^Ala^731^Arg^ (Fig. 5O, Fig. 5P), the H_2_-mediated promotion of Ca^2+^ influx was lost. These findings confirm that H_2_ relies on TRPC4AP to enhance cellular Ca^2+^ influx and that the binding site comprising 730^Arg^731^Arg^ on TRPC4-TRPC4AP at the CIRB motif is crucial for the opening of the Ca^2+^ channel.

### H_2_ enhanced cell motility by inducing Ca^2+^ influx to reconstruct cytoskeleton

Immunofluorescence (IF) analysis revealed predominant cytoplasmic localization of TRPC4AP in mesenchymal stem cells (MSCs) with no colocalization observed with the F-actin cytoskeletal network (Fig. 6A). Live-cell imaging over a 3-hour period demonstrated significantly accelerated F-actin cytoskeletal contraction dynamics in H₂-treated MSCs compared to untreated controls (Fig. 6B), indicative of enhanced cellular motility. Protein expression profiling showed coordinated upregulation of mesenchymal markers, with vimentin and α-smooth muscle actin (α-SMA) levels increasing post-H₂ treatment across multiple MSC subtypes (Fig. 6C, Fig. 6D). Ultrastructural analysis via transmission electron microscopy (TEM) revealed characteristic membrane-proximal aggregation of microfilament bundles following both acute (2-hour) and prolonged (24-hour) H₂ exposure (Fig. 6E).

**Fig. 6.**
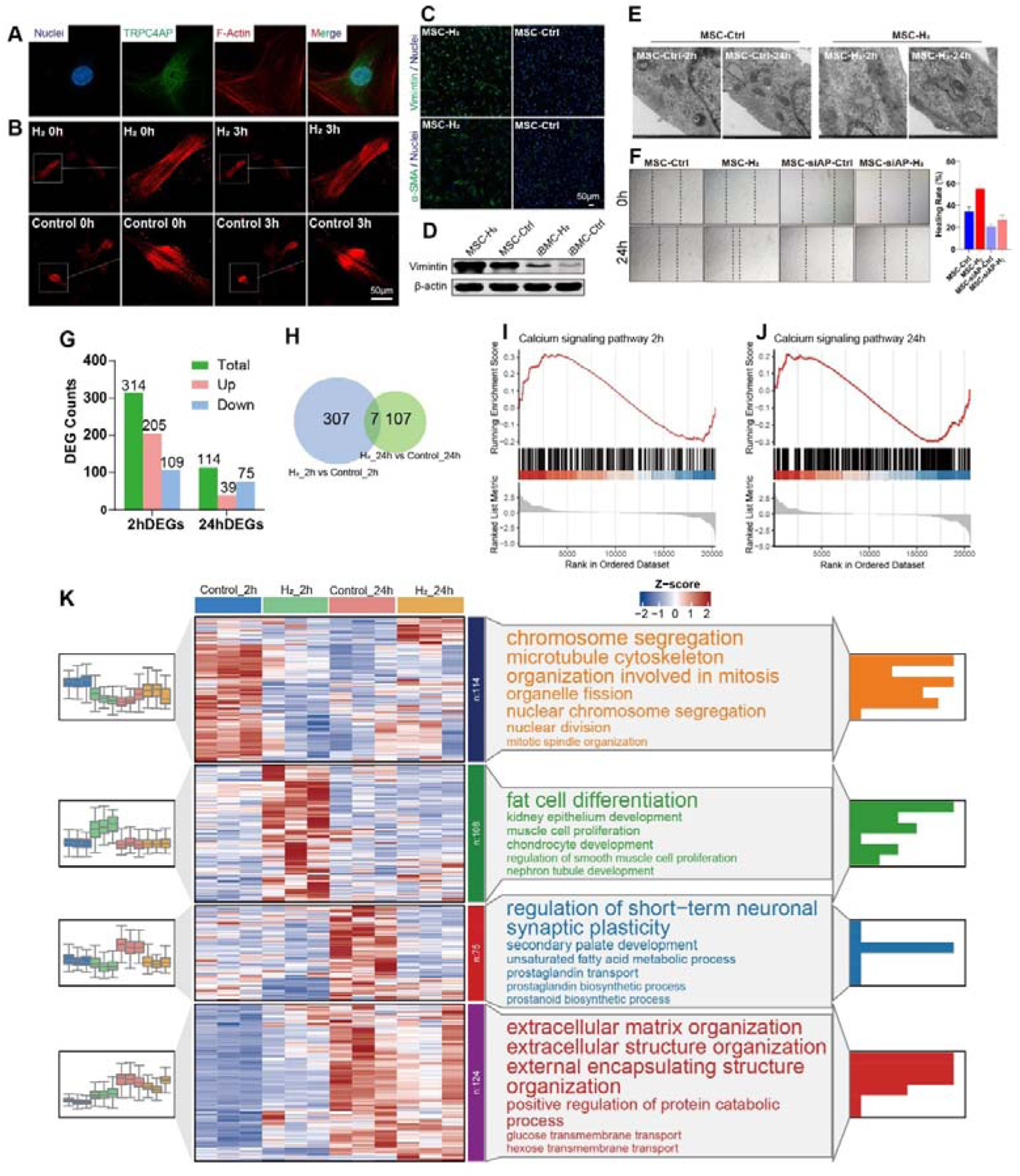
Multifaceted analysis of MSCs response to H_2_ treatment. A. Fluorescence co-localization results of F-Actin (red) and the TRPC4AP (green). B. Time-lapse F-Actin probes images of MSCs under H_2_-free (Control) and H_2_-medium. C. Immunofluorescence results of Vimentin and α-SMA in the Control and H_2_ groups of MSCs. D. Western Blot results of the effect of H_2_ on Vimintin expression in MSCs and iBMCs. E. Electron microscopy results of MSCs after treatment with H_2_ for 2 hours and 24 hours. F. Cell scratch results (left) and healing rate (right) for MSCs treated with H_2_ for 24 hours after siRNA AP treatment. G. Counts of differentially expressed genes (DEG) in MSCs after H_2_ treatment for 2 hours and 24 hours as revealed by RNA sequencing results. H. Venn diagram of differential genes between 2 hours and 24 hours post-treatment with H_2_. I&J. GSEA analysis of Calcium signaling pathway in MSCs treatment 2 hours (I) and 24 hours (J) under H_2_. K. Expression trends, heat maps, and significantly enriched Biological Process terms for each cluster of differentially expressed genes (DEGs) in MSCs; the left side depicts box plot diagrams illustrating the expression trends of DEGs across each cluster in Control and H_2_-treated samples; the center section presents heat maps of DEGs for each cluster, with color gradients from blue to red representing increasing expression levels; the right side shows the top 6 significantly enriched pathways from GO Biological Process enrichment analysis of DEGs in each cluster, sorted by p-value, along with the corresponding bar charts.

Functional assessments through scratch wound assays demonstrated significant H₂-mediated migration enhancement, with wound closure rates reaching 58% in treated groups versus 38% in controls at 24 hours. This pro-motility effect was substantially attenuated by trpc4ap knockdown, confirming pathway specificity. While histamine (a calcium agonist positive control) induced moderate calcium influx (Figure.S6A, Figure.S6B) and upregulated vimentin (Figure.S6C), α-SMA (Figure.S6D), and collagen I (Figure.S6D) expression, its efficacy remained inferior to H₂ stimulation.

Mechanistic investigation through time-course transcriptomics identified seven consensus differentially expressed genes (DEGs) common to both 2-hour and 24-hour H₂ treatments (Fig. 6G, Fig. 6H). Gene set enrichment analysis (GSEA) revealed significant calcium channel-related pathway activation at both time points (Fig.6I, Fig. 6J), corroborating the functional data.

These integrated findings establish that H₂-induced extracellular calcium influx primarily orchestrates cytoskeletal reorganization and motility enhancement in MSCs through TRPC4AP-dependent mechanisms.

RNA-seq analysis revealed that individual clustering showed upregulated genes in the MSCs-2h-H₂ group were primarily enriched in extracellular matrix (ECM) organization and tubule development (Figure. S7A), indicating significant modulations in the cytoskeleton. In the MSCs-24h-H₂ group, upregulated genes were mainly associated with muscle movement and Ca²⁺ transport (Figure. S7B), suggesting enhanced cell motility. Joint clustering analysis indicated that upregulated genes in the MSCs-2h-H₂ group were predominantly linked to fat cell, epithelium, and muscle cell differentiation, as well as muscle cell proliferation (Fig. 6K).

The effect of H_2_ was validated in the HUVEC cell line, demonstrating that H₂ similarly enhances the migratory capabilities of cells, evidenced by accelerated wound healing speed (Figure. S8A) and rapid reorganization of the cytoskeleton (Figure. S8B). Transcriptomic analysis conducted 2 hours post H₂ treatment (2h-H_2_ vs 2h-Ctrl) revealed significant inter-group differences, with an increased number of DEGs (Figure. S8C-S8E). GO and KEGG enrichment analyses indicated upregulation of processes related to development and ion regulation, along with cytoskeletal gene enrichment and notable activation of the calcium signaling pathway (Figure. S8F). This suggests that H₂ may modulate cytoskeletal remodeling in HUVECs through transient Ca²⁺ influx. By 24 hours, the upregulated genes were primarily associated with cytokine signaling, while genes related to Ca^2+^ influx were downregulated (Figure. S8F), indicating a shift towards Ca^2+^ efflux or cessation of influx, thereby initiating downstream signaling cascades.

### H_2_ Effects on Intracellular Cations

The TRP channel is a non-selective cation channel composed of four subunits, which is permeable to a variety of cations, including Ca^2+^, K⁺, and Na⁺ (Figure. S3). These cations exhibit competitive interactions within the channel, with Ca^2+^ generally being more permeable than Na⁺ and K⁺. In this study, we used Na⁺/K⁺ ion probes to investigate the ion influx in MSCs (Figure. S3A). We found that after adding H₂ to the regular culture medium, there was a slight increase in Na⁺ levels in the cytoplasm, while K⁺ influx remained unchanged. When the medium was switched to a Ca2+-free solution, Na⁺ influx continued to rise, while K⁺ influx still showed no significant change. These results suggest that hydrogen gas induces cation influx through TRPC4, where Ca^2+^ exhibits a much higher permeability than Na⁺ and K⁺. The slight increase in Na⁺ may be related to the NAX system, which helps expel excess Ca^2+^ that enters the cell. A similar phenomenon was observed in HUVECs (Figure. S3B).

## Discussion

Ca²⁺, as ubiquitous secondary messengers, orchestrate critical biological processes in eukaryotic cells, including gene expression, metabolic homeostasis, cell cycle progression, cellular motility, and neuromuscular signaling[1, 12, 28]. This study reveals that H₂ specifically activates the TRPC4-TRPC4AP signaling axis to induce calcium influx, uncovering for the first time the molecular mechanism by which H₂ functions as a novel calcium signaling agonist. To elucidate the molecular targets of H₂, we integrated computational structural biology with functional validation experiments: Molecular docking simulations identified a dual-arginine motif (730Arg-731Arg) within the carboxyl-terminal CIRB domain of TRPC4 as the H₂-binding pocket. Subsequent site-directed mutagenesis confirmed that this motif serves as a critical regulatory site for calcium channel activation in the TRPC4AP-TRPC4 complex. Molecular dynamics simulations further suggested that H₂ enhances the conformational stability of the ion channel by restructuring the hydrogen-bonding network of arginine residues and modulating local electrostatic potential. These findings not only establish a novel paradigm for gasotransmitter-mediated calcium homeostasis regulation but also provide an innovative structural framework for targeted drug design toward TRPC4 channels.

### Hydrogen Gas: Broad-Spectrum Effects and Calcium Homeostasis Regulation

According to the Human Protein Atlas[29], the TRPC4AP gene and its encoded protein are ubiquitously expressed across diverse tissue types (Figure. S9A, Figure. S9C). Although proteomic data for TRPC4 remain limited, transcriptomic profiling reveals tissue-specific enrichment of TRPC4 mRNA in the endometrium, seminal vesicles, and smooth muscle tissues (Figure. S9B, Figure. S9D). Strikingly, H₂-induced Ca²⁺ influx demonstrates broad-spectrum functionality, as evidenced by extracellular calcium entry triggered by hydrogen in mesenchymal stem cells, endothelial cells, epithelial cells, fibroblasts, osteoblasts, and cardiomyocytes (Fig.1A, Fig. 2A-F, Fig. 3J). Furthermore, in vivo experiments substantiate this universality: hydrogen exposure significantly amplifies Ca²⁺ transients in mouse primary motor cortex (M1) neurons (Fig. 2G-J), indicating H₂ enhances neuronal Ca²⁺ concentrations to potentiate synaptic transmission. These findings collectively validate the pan-cellular nature of H₂-mediated calcium signaling across both in vitro and in vivo systems.

Cellular viability critically depends on maintaining cytosolic Ca²⁺ within a narrow physiological range (∼200 nM in quiescent cells)[30]. Dysregulated calcium accumulation or depletion triggers pathological cascades leading to cellular senescence or apoptosis. This equilibrium is governed by a multi-tiered regulatory network: (1) PMCA (plasma membrane Ca²⁺-ATPase) and NCX (sodium-calcium exchanger)-mediated Ca²⁺ extrusion[17, 31]; (2) ER/SR calcium buffering and storage; (3) coordinated gating of plasma membrane calcium channels[32]. Our data reveal that H₂-induced Ca²⁺ elevation exhibits spatiotemporal compartmentalization: early-phase Ca²⁺ influx via TRPC4 channels is partially sequestered into the ER (Fig. 2G, Fig. 2H), while later-phase Ca²⁺ efflux occurs predominantly through NCX transporters (Fig.3J-L).

Notably, acute H₂ treatment elicits dual beneficial effects: (1) Cell viability remains uncompromised (Fig. 3A, Fig. 3G) with preserved mitochondrial integrity (Fig.3B-D)[33], confirming absence of cytotoxicity; (2) Further measurement of Ca-Mg-ATPase activity at different time points revealed that the enzyme activity in the H_2_ group increased over time, showing a gradually rising trend (Fig.3L), and reached more than six times that of the control group after 30 minutes. (3) Another channel for calcium extrusion at the cell membrane is NXC, and since the Na^+^ concentration did not change significantly (Figure S3B, D), the role of this channel in calcium extrusion is excluded.

### H₂-mediated Ca²⁺ influx originates extracellularly and depends on TRPC4 channel activity

TRPC ion channels constitute a class of non-selective cation channels that play a pivotal role in regulating a wide range of cellular functions, including nociception[34], mechanosensation[35], and signal transduction processes[36]. TRPC4 is broadly expressed across human tissues, with the Human Atlas Database identifying its presence in multiple organs, such as the heart, vasculature, brain, lungs, and kidneys (Figure.S9A, Figure.S9D). Functionally, TRPC4 can assemble as a homotetrameric channel or form heterotetrameric complexes with TRPC1 and TRPC5[32, 37]. Among these three subunits, TRPC4 and TRPC5 exhibit a higher degree of sequence homology. Accumulating research evidence suggests that TRPC1 functions as a negative regulator of TRPC4 and TRPC5, thereby modulating their physiological roles[38].

Through the use of calcium-free culture media replacement, pharmacological calcium depletion, and calcium ion tracing in both the mitochondria and endoplasmic reticulum[39], we determined that hydrogen-induced calcium influx primarily originates from extracellular sources (Fig. 1G, Fig. 1H), with minimal contributions from intracellular calcium stores, including the ER (Fig. 1K) and Mito (Fig 1L). Given these findings, we directed our focus towards Ca²⁺ channels located on the plasma membrane. Transcriptomic sequencing enabled us to quantify the expression levels of all proteins associated with calcium ion channels (Figure.S4A–S4C). Subsequent inhibitor studies and siRNA knockdown experiments further refined our focus to TRPC1, TRPC4, and TRPC5 as the key channels involved in this process.

Following the knockdown of TRPC1 and TRPC5, we observed that H₂ remained capable of inducing Ca²⁺ influx. However, while knockdown of TRPC1 resulted in a moderate attenuation of H₂-induced effects, silencing TRPC4 led to a significant reduction in Ca²⁺ influx. Additionally, transcriptomic sequencing revealed that among the TRPC family members, TRPC1 and TRPC4 exhibited higher baseline gene expression levels (Figure S4B), suggesting that TRPC1 may play a role in the assembly of the TRPC4 channel complex. Liu et al. demonstrated that TRPC4 is expressed in cerebral vasculature, where it is regulated by GPCRs and receptor tyrosine kinases, thereby facilitating Ca²⁺ influx and modulating vasodilation[40]. However, the effects of H₂ appear to be independent of Giα signaling (Figure. S4P) and may instead be mediated through TRPC4AP, which regulates the opening of TRPC4.

### TRPC4AP: A Molecular Switch for Hydrogen-Induced Activation of the TRPC4 Calcium Channel

TRPC4AP can directly bind to TRPC4 and interact with TRPC1, TRPC4, and TRPC5[26, 41]. To date, only the TRPC4-TRPC4AP complex has been demonstrated to trigger calcium loading upon depletion of ER Ca²⁺ stores, suggesting that this pathway may regulate Ca²⁺ influx[26]. Transcriptomic sequencing revealed that among TRPC-related genes, trpc4ap exhibited the highest baseline expression level (Figure S4B). Moreover, the effect of H₂ on Ca²⁺ influx was entirely dependent on TRPC4AP (Fig.4C, Fig.4D, Fig.4G, Fig.4H, Fig.4I, Fig.4J). Similarly, the influence of H₂ on cellular functions, including cytoskeletal dynamics (Figure 6B, E), migration (Fig. 6F), and differentiation (Fig.6C, Fig.6D), was strictly dependent on TRPC4AP expression. Furthermore, the effect of H₂ on Ca²⁺ influx was nearly abolished in TRPC4-deficient cells (Fig.4A, Fig.4B, Fig.4K, Fig.4L). These findings suggest that H₂ primarily relies on the molecular switch function of TRPC4AP to activate TRPC4 channels and facilitate Ca²⁺ influx.

### Elucidating the TRPC4-TRPC4AP Molecular Interaction Interface: Identification of Hydrogen (H**₂**)-Sensitive Functional Domains

The interaction between TRPC4 and TRPC4AP is currently unknown, with scant structural evidence available[42]. The IF staining indicates that TRPC4AP, resembling a scaffolding structure, is abundantly expressed throughout the cytoplasm (Fig. 6A). While the specific binding sites between TRPC4AP and TRP proteins remain unidentified. To address this, we performed molecular docking simulations between TRPC4 and TRPC4AP, which may relate to H_2_, predicting key binding sites at 730Arg and 731Arg on TRPC4 (Fig. 5E-5H). This binding site is located within the C-terminal CIRB functional box of TRPC4[43], a domain known to act as a switch for TRPC channels and previously identified as a binding site for IP3Rs, which regulate the endogenous SOCE, and calmodulin (CaM)[43], an inhibitor of IP3R, which binds to the CIRB box and activates TRPC4[44, 45] . The CIRB box is specifically present inTRPC4 subtypes and absent/different in TRPC1 and TRPC5, as shown by sequence alignment (Fig. 5A, Fig. 5B). This specificity suggests the unique role of this TRPC4 binding site within heterotetrameric complexes.

The predicted binding sites for TRPC4-TRPC4AP interaction, including ionic and hydrogen bond interactions, are primarily located within the basic amino acid-rich region of the C-terminal of TRPC4. Analysis of the frequency of amino acid occurrences in the 60 selected binding poses reveals that Arg and Lys are the most frequently represented amino acids. Notably, 730Arg and 731Arg in the CIRB-box, which appear as consecutive basic amino acids, show the highest frequency and are key sites critical to H_2_ function (Fig. 5C). Single and double amino acid mutations at the 730Arg and 730Arg/731Arg site and subsequent complementation indicated a critical dependency of H_2_ on the TRPC4-TRPC4AP binding site (Fig.7A). Additionally, Lys is found frequently, appearing 80 times in the 60 binding poses (Fig. 4F). This suggests that 715Lys and 716Lys, located in the Protein 4.1 module upstream of the CIRB motif, may also be potential sites affected by H_2_. Other possible sites include 737Arg and 738Arg within the CIRB, as well as 718Arg and 719Arg in the Protein 4.1 module (Fig.5B), all of which warrant further validation and investigation.

**Fig. 7.**
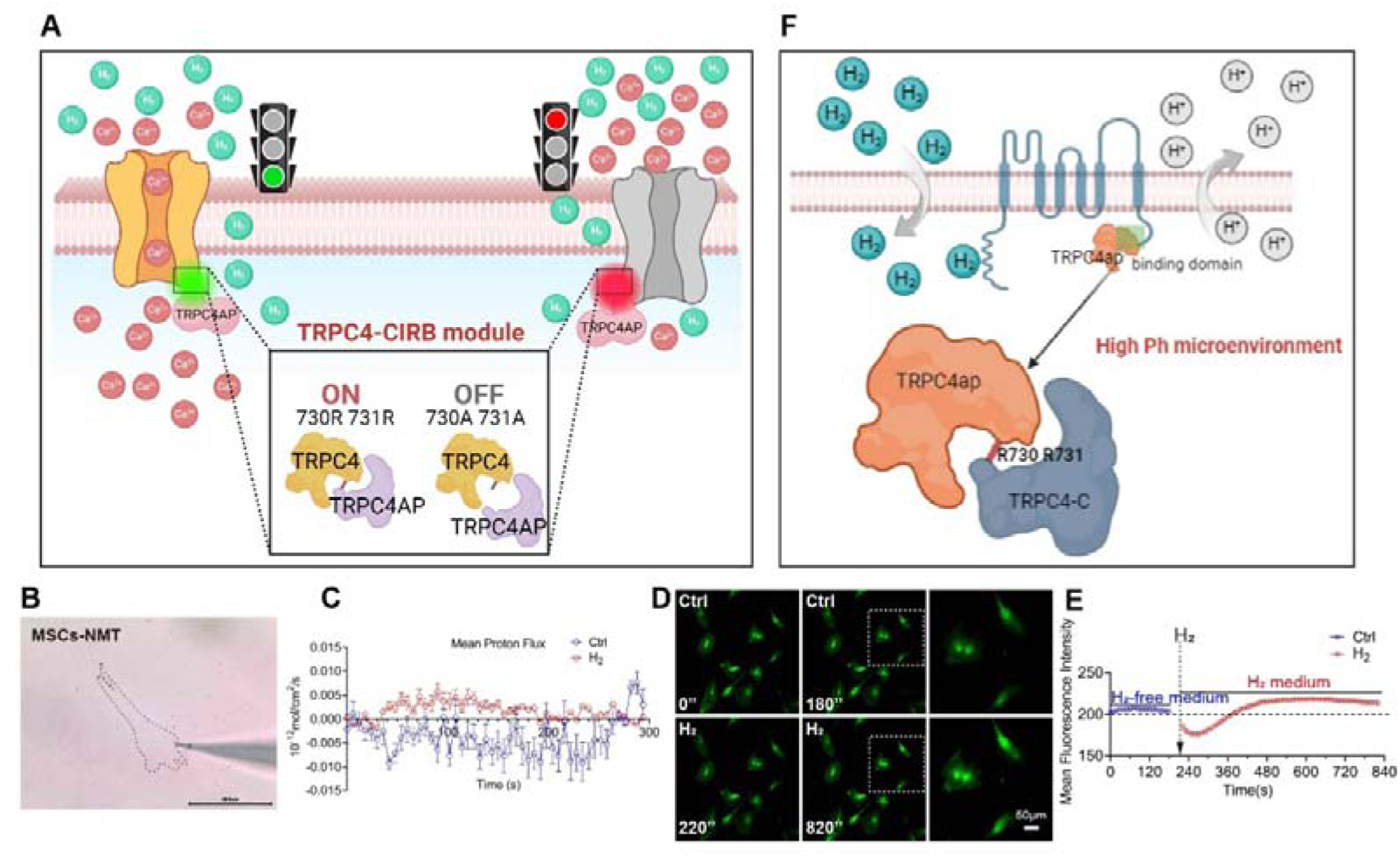
Potential molecular mechanism of H_2_ promoting [Ca^2+^]i by acting on the TRPC4-TRPC4ap binding site. A. Schematic representation of the H_2_-dependent binding site of TRPC4-TRPC4ap facilitating [Ca^2+^]i. B, Schematic diagram of live cell NMT detection of MSCs. C, Dynamic changes in proton flux across the plasma membrane of MSCs under the influence of H_2_. D & E, Time-lapse imaging showing the effect of H_2_ on intracellular pH changes and mean fluorescence intensity statistics.

The binding sites likely influenced by H_2_ are predominantly basic amino acids. In the exploratory extension of this study, NMT analysis indicates that H_2_ exposure leads to proton extrusion (Fig. 7B, Fig. 7C), while pH fluorescent probes suggest an increase in intracellular pH (Fig.7D, Fig.7E), creating an alkaline microenvironment[46, 47]. Such conditions are favorable for the stability of basic amino acids. These observations imply that H_2_ may indirectly modulate TRPC4-TRPC4AP binding by altering the intracellular acid-base environment. Previous studies have shown that eukaryotic plant plasma membranes possess hydrogenase activity, capable of decomposing and utilizing molecular H_2_, which in turn promotes proton extrusion and accelerates hypocotyl growth in mung beans , aligning with our findings. While hydrogenase activity has not yet been identified in animal cell membranes, our data suggest that H_2_ could alter protein interaction sites on the cell membrane by modulating local acidity or alkalinity. This points to a potential metabolic role for H_2_ at the cell membrane (Fig.7F), which requires further exploration.

### H_2_ May Regulate Multiple Biological Functions Through the Multifunctional Docking Site of TRPC4AP

In addition to its interaction with TRPC1/4/5, TRPC4AP exhibits multiple functions[48]. It has been identified as a key substrate-recognition component of the DCX

(DDB1-CUL4-X-box) E3 ubiquitin-protein ligase complex, which is crucial for regulating the cell cycle[49, 50] . Further investigations have revealed that the DCX (TRPC4AP) complex specifically mediates the polyubiquitination and subsequent degradation of MYC via the DesCEND (destruction via C-end degrons) pathway[50]. This mechanism also helps explain the typically lower expression levels of TRPC4AP observed in tumor cell lines compared to normal cells, resulting in distinct responses to H_2_, as shown in Figure .S10.

TRPC4AP, also known as TNF-R1 ubiquitous scaffolding and signaling protein, plays a pivotal role in the activation of NFκB1 upon TNFRSF1A ligation, potentially by bridging TNFRSF1A to the IKK signaling pathway[51]. Furthermore, TRPC4AP is instrumental in activating both NF-κ B and c-Jun N-terminal kinase (JNK) in response to TNF-α stimulation[52] . This multifaceted involvement highlights the essential role of TRPC4AP in the complex signaling networks that regulate cellular responses to TNF-α, demonstrating its importance in the orchestration of inflammatory and immune signaling pathways[53].

The multifaceted roles of TRPC4AP exhibit notable parallels with the functions of H_2_, particularly its ability to mitigate inflammatory storm responses and reduce inflammation-induced tissue damage[24, 54, 55]. Additionally, H_2_ has been shown to promote cellular proliferation and activation in various cell types, including epidermal stem cells, fibroblasts, and epithelial cells . Given that TRPC4AP is considered a key protein dependent on H_2_ for its function in facilitating Ca^2+^ influx, it is highly plausible that the diverse biological effects of H_2_ are linked to the ubiquitination and inflammation-related signaling pathways associated with TRPC4AP. This suggests that TRPC4AP may play a critical role in mediating the pleiotropic effects of H_2_ in cellular processes, particularly those involving inflammation and cellular activation.

### Therapeutic Implications and Future Directions

Endogenous H₂, predominantly produced by gut microbiota with a daily yield of 0.2-1.5 liters, serves as a critical modulator of metabolic processes[56, 57]. Within the intestinal ecosystem, H₂ exhibits dual functionality—participating in both its own biosynthesis and catabolism—thereby regulating secondary metabolite conversion. We propose that interindividual variations in gut microbiota composition drive heterogeneous H₂ levels, which may differentially influence Ca²⁺ flux dynamics and downstream physiological outcomes.

Excessive Ca²⁺ influx acts as a pathogenic driver across multiple cellular processes: Neurodegeneration: Mitochondrial calcium signaling disruption correlates with Alzheimer’s[58], Parkinson’s[59], and Huntington’s pathologies[60]. Immune Dysfunction: Calcium signaling pathways are indispensable for lymphocyte activation and immune disorder progression. While H₂ demonstrates unique calcium-modulating capacity, its therapeutic application requires stringent optimization of: dosage thresholds and temporal dynamics.

Our discovery of the H₂-TRPC4/TRPC4AP regulatory axis unveils a groundbreaking therapeutic approach for calcium homeostasis disorders. This molecular switch enables: (1) precision-targeted correction of calcium overload while preserving physiological fluctuations, (2) tissue-selective modulation of calcium-dependent signaling cascades, and (3) prevention of compensatory stress responses typically associated with conventional calcium channel blockers. These findings are anticipated to systematically elucidate the multifaceted physiological roles of H₂ in critical biological processes.

## Conclusion

This study elucidates the molecular mechanism by which H₂ functions as a first-in-class calcium signaling agonist, demonstrating its capacity to modulate calcium dynamics through the TRPC4-TRPC4AP channel axis. Our findings reveal that H₂ binds to a discrete arginine-rich motif (730Arg-731Arg) within the CIRB domain of TRPC4, triggering conformational changes that potentiate extracellular Ca²⁺ influx while maintaining calcium homeostasis. This gasotransmitter-like activity establishes H₂ as a unique endogenous regulator of calcium signaling networks.

## Author Contributions

Conceptualization P.Z. and X.M.; Data curation, P.Z., H.L., and Z.C.; Formal analysis, X.Z., X.W., Z.L., and S.J.; Funding acquisition, X.M. and P.Z.; Methodology, Z.C., P.Z., H.L., X.Z., Z.D., X.J., and J.W.; Visualization, P.Z., H.L., Z.C., and X.Z.; Writing – original draft, P.Z.; Writing – review & editing, X.Z., M.L., F.X. and X.M. All authors have read and agreed to the published version of the manuscript.

## Data availability statement

The original contributions presented in this study are included in the article/Supplementary.

## Acknowledgments

We thank Biorender (www.biorender.com) for their drawing assistance during the writing of this article.

## Ethics statement

The study was conducted in accordance with the Declaration of Helsinki.

## Supplemental Figs

**Figure S1.**
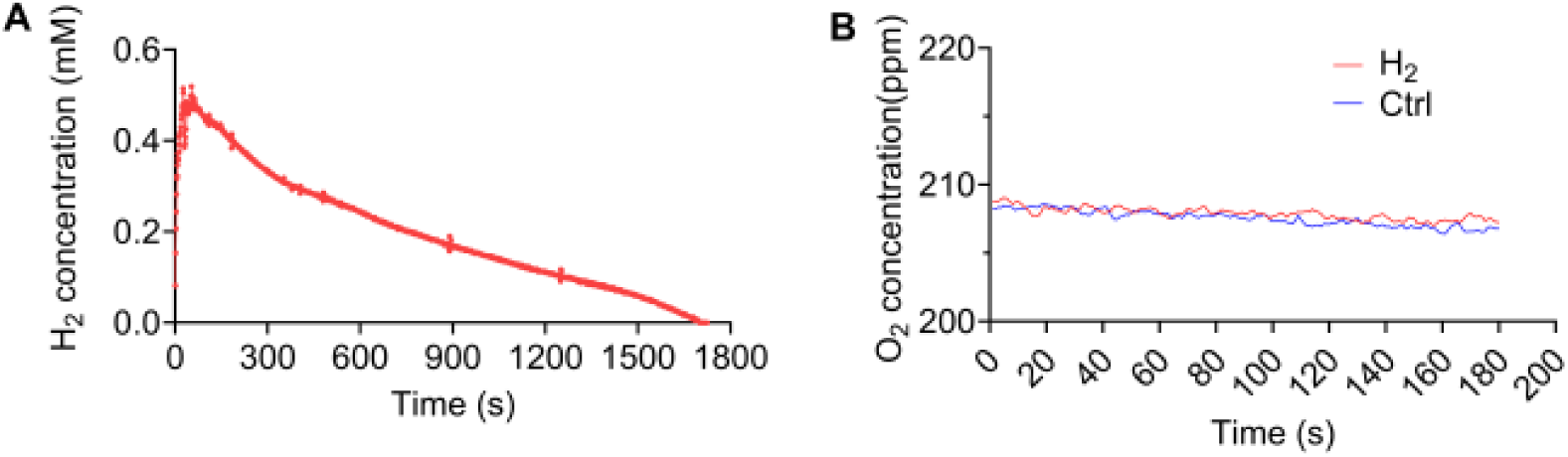
Real-time monitoring of H_2_ and O_2_ levels in the culture medium after one single administration of H_2_. (A) Real-time monitoring curve of H_2_ concentration. (B) Real-time monitoring curve of O_2_ concentration.

**Figure S2.**
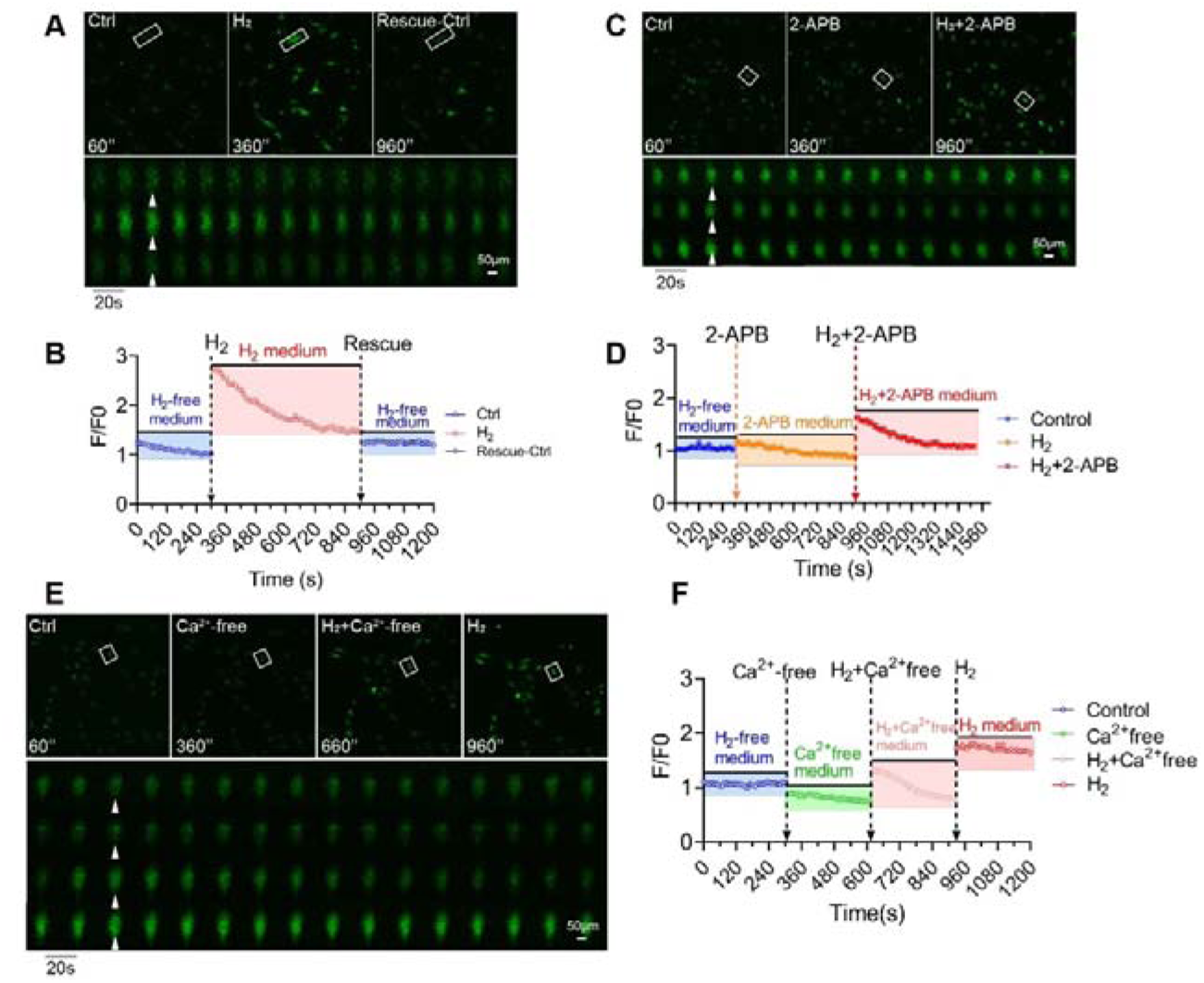
H_2_ enhances extracellular [Ca^2+^]i in HUVEC cells. (A) & (B) Time-lapse (A) images and the Fluo 4 averaged ΔF/F0 trace (B) of the [Ca^2+^]i changes under H_2_-free, H_2_-medium, and then back to H_2_-free conditions in HUVEC. (C) & (D) Time-lapse images (C) and the Fluo4 averaged F/F0 trace (D) of the [Ca^2+^]i changes under H_2_-free, 2-APB-medium, and then back to 2-APB+H_2_-medium in HUVEC. (E) & (F) Time-lapse images (E) and the Fluo4 averaged F/F0 trace of the [Ca^2+^]i changes under H_2_-free, Ca^2+^free-medium, H_2_+Ca^2+^free-medium, and then back to H_2_-medium conditions. White arrows in A, C, and E indicate cells at distinct time points within the time-lapse capture. Sampling rate, 10 sec.

**Figure S3.**
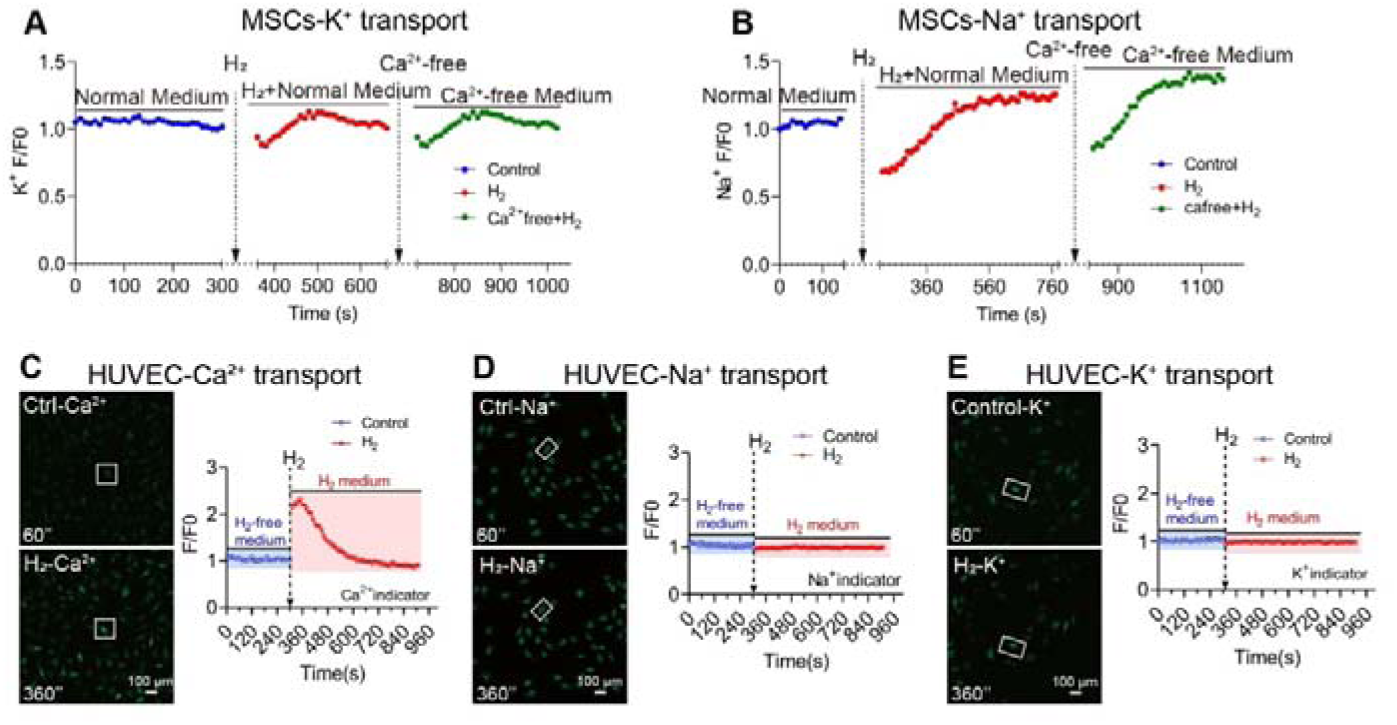
Time-lapse averaged ΔF/F0 trace for the influence of H_2_ on the influx of different cations in MSCs and HUVEC. (A) & (B) H_2_-induced K+ and Na+ transport under normal-medium, H_2_-medium and Ca^2+^-free medium conditions in MSCs. (C) (D) & (E) Comparison of the effects of H_2_ on Ca^2+^, K+, and Na+ influx in HUVEC cells.

**Figure S4.**
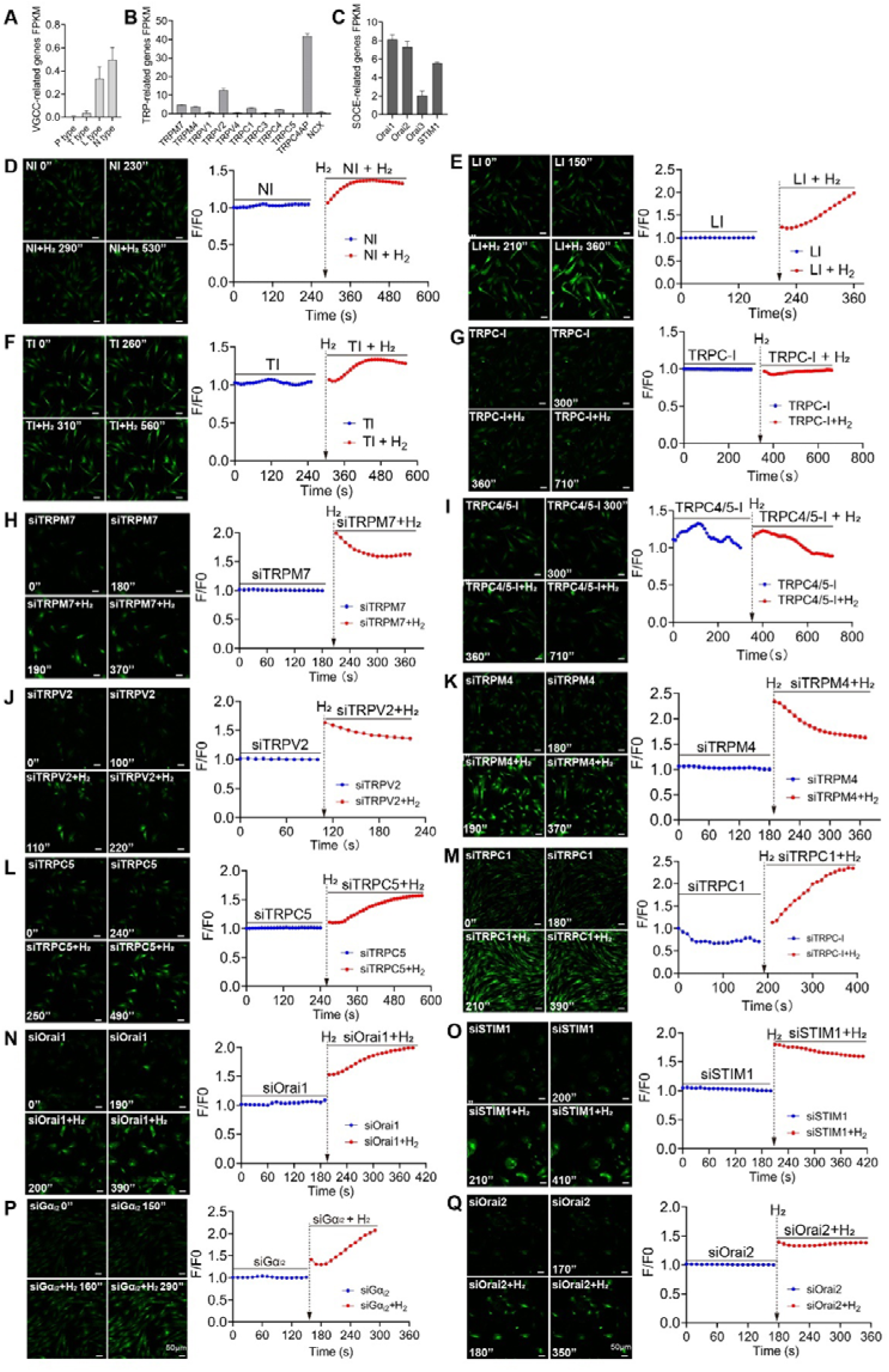
Screening for Ca^2+^ channels potentially dependent on H_2_ in MSCs using siRNA or channel inhibitors. (A)-(C) Bulk RNAseq analysis of MSCs reveals a comparative assessment of FPKM values for various types of Ca^2+^ channels and their associated genes expression. (D)-(F) &( I) Screening for Ca^2+^ channels potentially targeted by H_2_, including N-type, L-type, T-type, TRPC, and TRPC4/5, using different inhibitors. (H)-(J) & (Q) Screening for the potential influences of siRNA-mediated TRPM7, TRPV2, TRPM4, TRPC5, TRPC1, Orai1, STIM1, Orai2, and Gα_i2_ and TRPC4ap knockdown in the [Ca^2+^]i triggered by H_2_.

**Figure S5.**
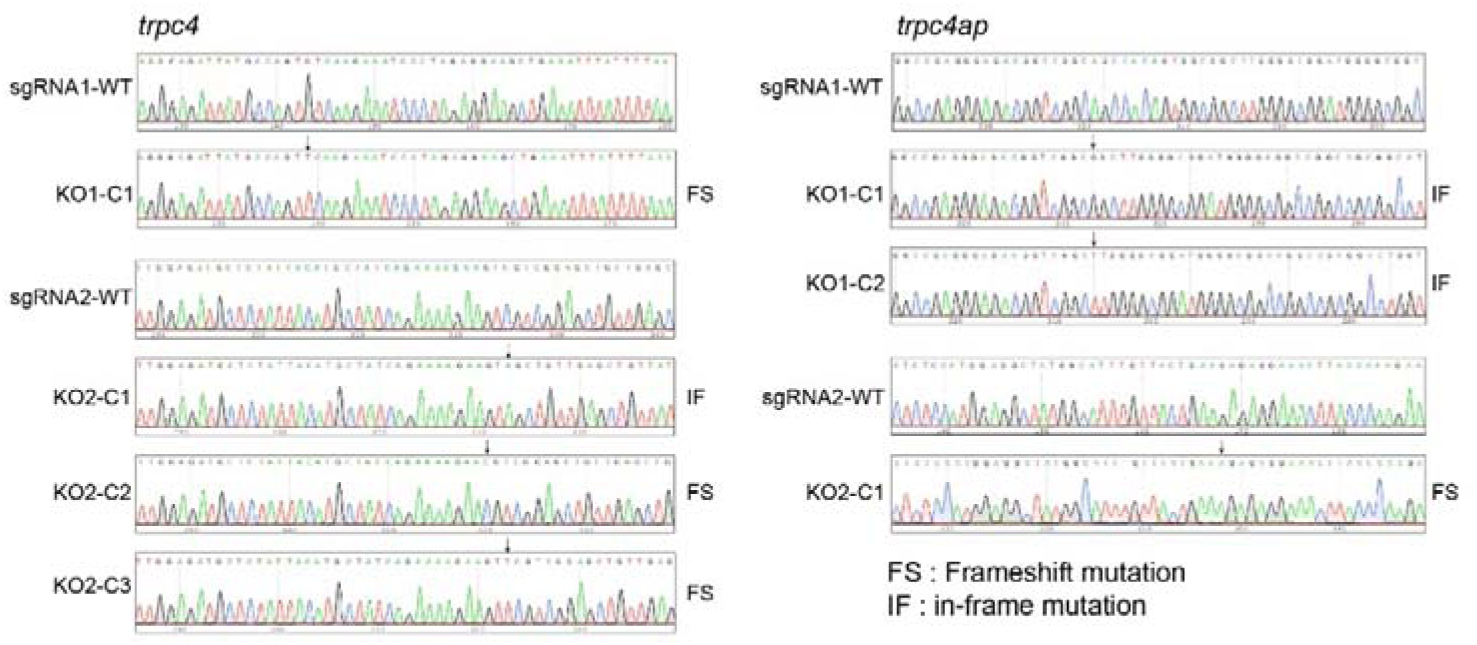
Validation of TRPC4 and TRPC4ap gene knockdown in monoclonal 293T cell lines. The mutant site is indicated by an arrow. Wild-type (WT) sequences of the corresponding exons were determined by Sanger sequencing and are presented with chromatograms. For each detected sequence variant, the detection frequency and the type of mutation are shown.

**Figure S6.**
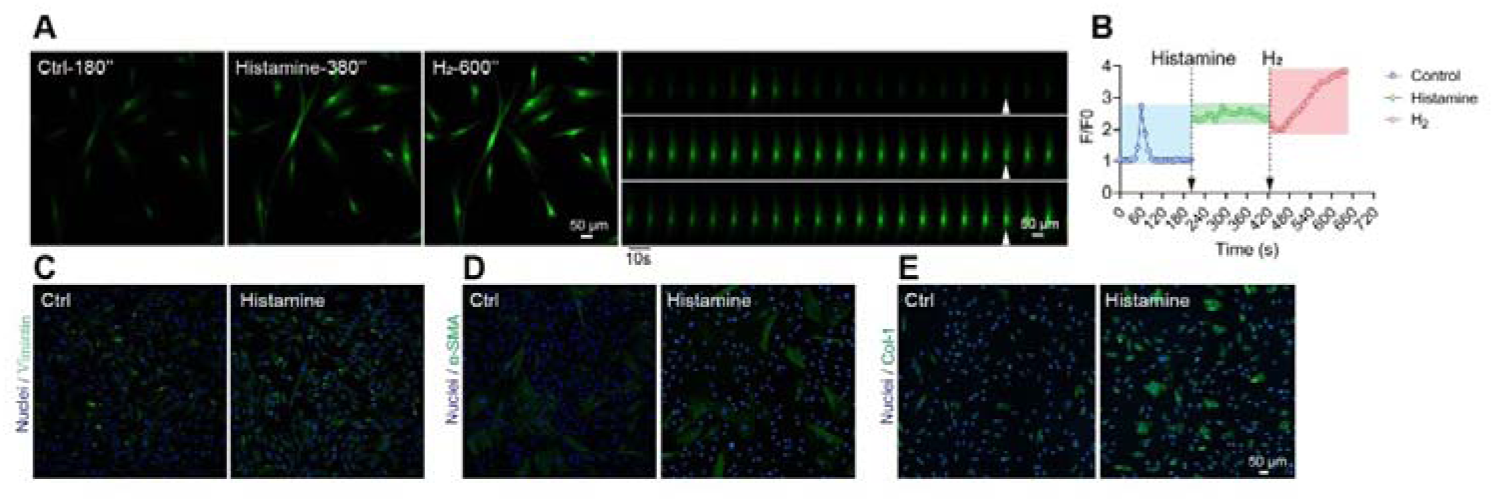
Comparative analysis of the effects of histamine, a Ca^2+^ agonist, and H_2_ on [Ca^2+^]i in MSCs, as well as their impact of histamine on the cytoskeleton. (A) & (B) Time-lapse (A) images and the Fluo4 averaged ΔF/F0 trace (B) of the [Ca^2+^]i changes under H_2_-free, Histamine, and then H_2_ conditions in MSCs. (C)-(E) Immunofluorescence results of Vimentin, α-SMA, and Collagen-1 in the Control and Histamine groups of MSCs.

## References

1. Clapham DE: Calcium signaling. Cell 2007, 131(6):1047–1058.

2. Zheng J, Zeng X, Wang S: Calcium ion as cellular messenger. In., vol. 58: Springer; 2015: 1–5.

3. Ghosh A, Ginty DD, Bading H, Greenberg ME: Calcium regulation of gene expression in neuronal cells. Journal of neurobiology 1994, 25(3):294–303.

4. Kahl CR, Means AR: Regulation of cell cycle progression by calcium/calmodulin-dependent pathways. Endocrine reviews 2003, 24(6):719–736.

5. Zheng JQ, Poo M-m: Calcium signaling in neuronal motility. Annu Rev Cell Dev Biol 2007, 23(1):375–404.

6. Kondratskyi A, Kondratska K, Skryma R, Klionsky DJ, Prevarskaya N: Ion channels in the regulation of autophagy. Autophagy 2018, 14(1):3–21.

7. Honda H, Kondo T, Zhao Q-L, Feril Jr LB, Kitagawa H: Role of intracellular calcium ions and reactive oxygen species in apoptosis induced by ultrasound. Ultrasound in medicine & biology 2004, 30(5):683–692.

8. Dolphin AC, Lee A: Presynaptic calcium channels: specialized control of synaptic neurotransmitter release. Nature Reviews Neuroscience 2020, 21(4):213–229.

9. Li HF, Li XY, Sun YQ, Zhi ZY, Song LF, Li M, Feng YM, Zhang ZH, Liu YF, Chen YJ: The Role of Ion Channels in Pulmonary Hypertension: A Review. Pulmonary Circulation 2025, 15(1):e70050.

10. Suzuki K: Thrombomodulin: A key regulator of intravascular blood coagulation, fibrinolysis, and inflammation, and a treatment for disseminated intravascular coagulation. Proceedings of the Japan Academy, Series B 2025, 101(2):75–97.

11. Makiyama F, Kawase S, Omi AW, Tanikawa Y, Kotani T, Shirayama T, Nishimura N, Kurihara T, Saito N, Takahashi J: Differential effects of structurally different lysophosphatidylethanolamine species on proliferation and differentiation in pre-osteoblast MC3T3-E1 cells. Scientific Reports 2025, 15(1):466.

12. Berridge MJ, Bootman MD, Roderick HL: Calcium signalling: dynamics, homeostasis and remodelling. Nature reviews Molecular cell biology 2003, 4(7):517–529.

13. Cartes-Saavedra B, Ghosh A, Hajnóczky G: The roles of mitochondria in global and local intracellular calcium signalling. Nature Reviews Molecular Cell Biology 2025:1–20.

14. Rizzuto R, Pinton P, Carrington W, Fay FS, Fogarty KE, Lifshitz LM, Tuft RA, Pozzan T: Close contacts with the endoplasmic reticulum as determinants of mitochondrial Ca^2+^ responses. Science 1998, 280(5370):1763–1766.

15. Hoth M, Fanger CM, Lewis RS: Mitochondrial regulation of store-operated calcium signaling in T lymphocytes. The Journal of cell biology 1997, 137(3):633–648.

16. Suzuki J, Kanemaru K, Ishii K, Ohkura M, Okubo Y, Iino M: Imaging intraorganellar Ca^2+^ at subcellular resolution using CEPIA. Nature communications 2014, 5(1):4153.

17. Raffaello A, Mammucari C, Gherardi G, Rizzuto R: Calcium at the center of cell signaling: interplay between endoplasmic reticulum, mitochondria, and lysosomes. Trends in biochemical sciences 2016, 41(12):1035–1049.

18. Emrich SM, Yoast RE, Trebak M: Physiological functions of CRAC channels. Annual Review of Physiology 2022, 84(1):355–379.

19. Nilius B, Owsianik G: The transient receptor potential family of ion channels. Genome biology 2011, 12:1–11.

20. Bacsa B, Tiapko O, Stockner T, Groschner K: Mechanisms and significance of Ca^2+^ entry through TRPC channels. Current opinion in physiology 2020, 17:25–33.

21. Zhang M, Ma Y, Ye X, Zhang N, Pan L, Wang B: TRP (transient receptor potential) ion channel family: structures, biological functions and therapeutic interventions for diseases. Signal Transduction and Targeted Therapy 2023, 8(1):261.

22. Bechem M, Schramm M: Calcium-agonists. Journal of molecular and cellular cardiology 1987, 19:63–75.

23. Tarr TB, Malick W, Liang M, Valdomir G, Frasso M, Lacomis D, Reddel SW, Garcia-Ocano A, Wipf P, Meriney SD: Evaluation of a novel calcium channel agonist for therapeutic potential in Lambert–Eaton myasthenic syndrome. Journal of Neuroscience 2013, 33(25):10559–10567.

24. Ohta S: Molecular hydrogen as a preventive and therapeutic medical gas: initiation, development and potential of hydrogen medicine. Pharmacology & therapeutics 2014, 144(1):1–11.

25. Ohsawa I, Ishikawa M, Takahashi K, Watanabe M, Nishimaki K, Yamagata K, Katsura K-i, Katayama Y, Asoh S, Ohta S: Hydrogen acts as a therapeutic antioxidant by selectively reducing cytotoxic oxygen radicals. Nature medicine 2007, 13(6):688–694.

26. Mace KE, Lussier MP, Boulay G, Terry-Powers JL, Parfrey H, Perraud AL, Riches DW: TRUSS, TNF-R1, and TRPC ion channels synergistically reverse endoplasmic reticulum Ca^2+^ storage reduction in response to m1 muscarinic acetylcholine receptor signaling. Journal of cellular physiology 2010, 225(2):444–453.

27. Abramson J, Adler J, Dunger J, Evans R, Green T, Pritzel A, Ronneberger O, Willmore L, Ballard AJ, Bambrick J: Accurate structure prediction of biomolecular interactions with AlphaFold 3. Nature 2024, 630(8016):493–500.

28. Staruschenko A, Alexander RT, Caplan MJ, Ilatovskaya DV: Calcium signalling and transport in the kidney. Nature Reviews Nephrology 2024, 20(8):541–555.

29. Pontén F, Jirström K, Uhlen M: The Human Protein Atlas—a tool for pathology. The Journal of Pathology: A Journal of the Pathological Society of Great Britain and Ireland 2008, 216(4):387–393.

30. Peacock M: Calcium metabolism in health and disease. Clinical Journal of the American society of nephrology 2010, 5(Supplement_1):S23–S30.

31. Di Leva F, Domi T, Fedrizzi L, Lim D, Carafoli E: The plasma membrane Ca^2+^ ATPase of animal cells: structure, function and regulation. Archives of biochemistry and biophysics 2008, 476(1):65–74.

32. Brini M, Calì T, Ottolini D, Carafoli E: The plasma membrane calcium pump in health and disease. The FEBS journal 2013, 280(21):5385–5397.

33. Zhao P, Dang Z, Liu M, Guo D, Luo R, Zhang M, Xie F, Zhang X, Wang Y, Pan S: Molecular hydrogen promotes wound healing by inducing early epidermal stem cell proliferation and extracellular matrix deposition. Inflammation and Regeneration 2023, 43(1):22.

34. Sun Z-C, Ma S-B, Chu W-G, Jia D, Luo C: Canonical transient receptor potential (TRPC) channels in nociception and pathological pain. Neural Plasticity 2020, 2020(1):3764193.

35. Eijkelkamp N, Quick K, Wood JN: Transient receptor potential channels and mechanosensation. Annual review of neuroscience 2013, 36(1):519–546.

36. Pedersen SF, Owsianik G, Nilius B: TRP channels: an overview. Cell calcium 2005, 38(3-4):233–252.

37. Duan J, Li J, Zeng B, Chen G-L, Peng X, Zhang Y, Wang J, Clapham DE, Li Z, Zhang J: Structure of the mouse TRPC4 ion channel. Nature Communications 2018, 9(1):3102.

38. Kim J, Ko J, Myeong J, Kwak M, Hong C, So I: TRPC1 as a negative regulator for TRPC4 and TRPC5 channels. Pflügers Archiv-European Journal of Physiology 2019, 471:1045–1053.

39. Malli R, Frieden M, Trenker M, Graier WF: The role of mitochondria for Ca2+ refilling of the endoplasmic reticulum. Journal of Biological Chemistry 2005, 280(13):12114–12122.

40. Liu D, Xiong S, Zhu Z: Imbalance and dysfunction of transient receptor potential channels contribute to the pathogenesis of hypertension. Science China Life Sciences 2014, 57:818–825.

41. Mace KE: Intersecton of cellular signaling cascades with TRPC ion channel responses and calcium ion homeostasis. University of Colorado Health Sciences Center; 2009.

42. Poduslo SE, Huang R, Huang J, Smith S: Genome screen of late-onset Alzheimer’s extended pedigrees identifies TRPC4AP by haplotype analysis. American Journal of Medical Genetics Part B: Neuropsychiatric Genetics 2009, 150(1):50–55.

43. Zhu MX: Multiple roles of calmodulin and other Ca 2+-binding proteins in the functional regulation of TRP channels. Pflügers Archiv 2005, 451:105–115.

44. Freichel M, Tsvilovskyy V, Camacho-Londoño JE: TRPC4-and TRPC4-containing channels. Mammalian Transient Receptor Potential (TRP) Cation Channels: Volume I 2014:85–128.

45. Vinayagam D, Quentin D, Yu-Strzelczyk J, Sitsel O, Merino F, Stabrin M, Hofnagel O, Yu M, Ledeboer MW, Nagel G: Structural basis of TRPC4 regulation by calmodulin and pharmacological agents. Elife 2020, 9:e60603.

46. Zhang X, Xie F, Zhang Z, Adzavon YM, Su Z, Zhao Q, LeBaron TW, Li Q, Lyu B, Liu G: Hydrogen evolution and absorption phenomena in the plasma membrane of Vigna radiata and Capsicum annuum. Journal of Plant Growth Regulation 2023, 42(1):249–259.

47. Han M, Yang H, Yu G, Jiang P, You S, Zhang L, Lin H, Liu J, Shu Y: Application of Non-invasive Micro-test Technology (NMT) in environmental fields: A comprehensive review. Ecotoxicology and Environmental Safety 2022, 240:113706.

48. Zeng C, Tian F, Xiao B: TRPC channels: prominent candidates of underlying mechanism in neuropsychiatric diseases. Molecular neurobiology 2016, 53(1):631–647.

49. Choi SH, Wright JB, Gerber SA, Cole MD: Myc protein is stabilized by suppression of a novel E3 ligase complex in cancer cells. Genes & development 2010, 24(12):1236–1241.

50. Koren I, Timms RT, Kula T, Xu Q, Li MZ, Elledge SJ: The eukaryotic proteome is shaped by E3 ubiquitin ligases targeting C-terminal degrons. Cell 2018, 173(7):1622–1635. e1614.

51. Terry Powers JL, Mace KE, Parfrey H, Lee S-J, Zhang G, Riches DW: TNF receptor-1 (TNF-R1) ubiquitous scaffolding and signaling protein interacts with TNF-R1 and TRAF2 via an N-terminal docking interface. Biochemistry 2010, 49(36):7821–7829.

52. Soond SM, Terry JL, Colbert JD, Riches DW: TRUSS, a novel tumor necrosis factor receptor 1 scaffolding protein that mediates activation of the transcription factor NF-κB. Molecular and cellular biology 2003, 23(22):8334–8344.

53. Soond SM, Terry JL, Riches DW: TRUSS, a tumor necrosis factor receptor-1-interacting protein, activates c-Jun NH_2_-terminal kinase and transcription factor AP-1. FEBS letters 2006, 580(19):4591–4596.

54. Tian Y, Zhang Y, Wang Y, Chen Y, Fan W, Zhou J, Qiao J, Wei Y: Hydrogen, a novel therapeutic molecule, regulates oxidative stress, inflammation, and apoptosis. Frontiers in physiology 2021, 12:789507.

55. Yang M, Dong Y, He Q, Zhu P, Zhuang Q, Shen J, Zhang X, Zhao M: Hydrogen: a novel option in human disease treatment. Oxidative Medicine and Cellular Longevity 2020, 2020(1):8384742.

56. Zhang Y, Xu J, Yang H: Hydrogen: An endogenous regulator of liver homeostasis. Frontiers in Pharmacology 2020, 11:877.

57. Tadross MR, Dick IE, Yue DT: Mechanism of local and global Ca^2+^ sensing by calmodulin in complex with a Ca^2+^ channel. Cell 2008, 133(7):1228–1240.

58. Rahman MH, Bajgai J, Fadriquela A, Sharma S, Trinh Thi T, Akter R, Goh SH, Kim C-S, Lee K-J: Redox effects of molecular hydrogen and its therapeutic efficacy in the treatment of neurodegenerative diseases. Processes 2021, 9(2):308.

59. Ostojic SM: Inadequate production of H_2_ by gut microbiota and Parkinson disease. Trends in Endocrinology & Metabolism 2018, 29(5):286–288.

60. Paul BD: Cysteine metabolism and hydrogen sulfide signaling in Huntington’s disease. Free Radical Biology and Medicine 2022, 186:93–98.

