## supplemental materials for "Hydrogen-Induced Calcium Influx via the TRPC4-TRPC4AP Axis"

*** These authors share first authorship**

**TABLE**

| **RAGENT or RESOURSE** | **SOURCE** | **IDENTIFIER** |
| --- | --- | --- |
| **Antibodies** |  |  |
| Rabbit anti-TRPC4AP | Proteintech | 29310-1-AP |
| Mouse anti-αSMA | Cell SignalingTechnology | 19245 |
| Rabbit anti-Vimentin | Cell SignalingTechnology | 5741 |
| Mouse anti-Beta actin | Proteintech | 66009-1-1g |
| Rabbit anti- CollagenⅠ | Abcam | AB254113 |
| secondary antibodies (Alexa Fluor 488) | Cell SignalingTechnology | 4412 |
| **Chemicals** |  |  |
| Ca^2+^-GPCR,Fluo-4,AM (5μM,for Live-cell) | KeyGEN | KGAF024 |
| Na^+^-ENG-2,AM, (5 μM, for Live-cell) | Maokang | MX4514 |
| K^+^-EPG-4,AM, (5 μM , for Live-cell) | Maokang | MX4521 |
| DAPI | ThermoFisher | R37606 |
| ER-tracker ( 1 μM ) | YEASEN | 40764ES20 |
| Mito-tracte r(100 nM) | YEASEN | 40741ES50 |
| Stem cell culture-medium | iCell | PriMed-icell-012-sf |
| JC-1 (2 μM ) | ThermoFisher | T3168 |
| CCK-8 | Beyotime | C0038 |
| Trypsin EDTA(0.25%) | ThermoFisher | 25200056 |
| DMEM high | Gibco | 11865092 |
| 1640 culture-medium | Gibco | 11875119 |
| Ca^2+^-free medium | iCell | PriMed-icell-012-sf |
| Phosphate buffered saline,PH=7.4 | Solarbio | P1003-2L |
| LipofectamineTM 2000 Transfection Reagent | ThermoFisher | 11668019 |
| Triton TM X-100 | Solarbio | T8200 |
| CoraLite®594 F-Actin | Proteintech | PF00003 |
| 4% paraformaldehyde (PFA) | ThermoFisher | J61899.AP |
| Bovine Serum Albumin | Gibco | A5256701 |
| BCA protein concentration determination kit | Beyotime | P0010 |
| TRIzol^TM^ Reagent | ThermoFisher | 15596018CN |
| OPTI-MEM | Gibco | 2898884 |
| SKF-96365 hydrochloride(10μM) | MedChemExpress | HY-100001 |
| ML204(0.5μM) | MedChemExpress | HY-12949 |
| Mibefradil dihydrochloride T(2.7μM) | MedChemExpress | HY-15553A |
| Amlodipine L(30μM) | MedChemExpress | HY-B0317 |
| PD173212 N (10μM) | MedChemExpress | HY-103318 |
| Histamine (10μM) | MedChemExpress | HY-B1204 |
| 2-APB(100μM) | MedChemExpress | HY-W009724 |
| Penicillin-Streptomycin | Gibco | 15070063 |
| Total Protein Extraction (TPE(TM) ) | Sangon | C006225 |
| Ca^2+^-Rhod-2, AM, (5μM,for Live-cell) | YEASEN | 40776ES72 |
| Ionomycin calcium (2μM) | MedChemExpress | HY-13434A |
| Ca++Mg++-ATPase Assay Kit | BIOSS | AK266 |
| Hank's Balanced Salt Solution | ThermoFisher | 14175095 |
| Total Protein Extraction | Sangon | C006225 |
| NucGreen™ Dead 488 ReadyProbes™ | ThermoFisher | S7020 |
| **Experimental models: Cell lines** |  |  |
| C57 | Vitalriver | N/A |
| MSCs | BeijingMaternityHospital | N/A |
| MC3TE-e1 | Peking Union Cell Bank | N/A |
| C2C12 | Peking Union Cell Bank | N/A |
| iBMSCs | icell | N/A |
| NIH-3T3 | Peking Union Cell Bank | N/A |
| ESF | Peking Union Cell Bank | N/A |
| PC12 | Peking Union Cell Bank | N/A |
| HUVEC | Peking Union Cell Bank | N/A |
| 293T | Peking Union Cell Bank | N/A |
| PX459 | Peking Union Cell Bank | N/A |
| **SiRNASequence (5’-3’)** |  |  |
| si-TRPC4AP-homo  GAGGAAACUUACCCAAGAATT | This Study | Gene Pharma |
| si-TRPM7-homo  AACAUCAGACGAACAGAAUUAGUUG | This Study | Gene Pharma |
| si-TRPM4-homo  AUGUGUAACUGAACAUGGCTT | This Study | Gene Pharma |
| si-TRPV2-homo  GCUUCCUUCUGAUCUACUUTT-3 | This Study | Gene Pharma |
| si-ORAI1-homo  CGUGCACAAUCUCAACUCGTT | This Study | Gene Pharma |
| si-ORAI2-homo  AACCGUUUGGUUCAAUGAGG | This Study | Gene Pharma |
| si-STIM1-homo  GGCUCUGGAUACAGUGCUCTT | This Study | Gene Pharma |
| si-TRPC1-homo  GCGACAAGGGUGACUAUUATT | This Study | Gene Pharma |
| si-TRPC4-homo  GGUCAGACUUGAACAGGCATT | This Study | Gene Pharma |
| si-TRPC5-homo  UACCAAGGUAUAGAAAUGACUUGGG | This Study | Gene Pharma |
| **Others** |  |  |
| 96-well cell culture plate | Corning | 3548 |
| 6-well cell culture plate | Corning | 3335 |
| Nunc™ EasYFlask™ Cell Culture Flasks-T25 | ThermoFisher | 156367 |
| Nunc™ EasYFlask™ Cell Culture Flasks-T75 | ThermoFisher | 156499 |
| Glass bottom cell culture dish (20mm) | NEST | 801001 |
| Centrifuge tube-15ml | NEST | 601001 |
| Centrifuge tube-50ml- | NEST | 602002 |

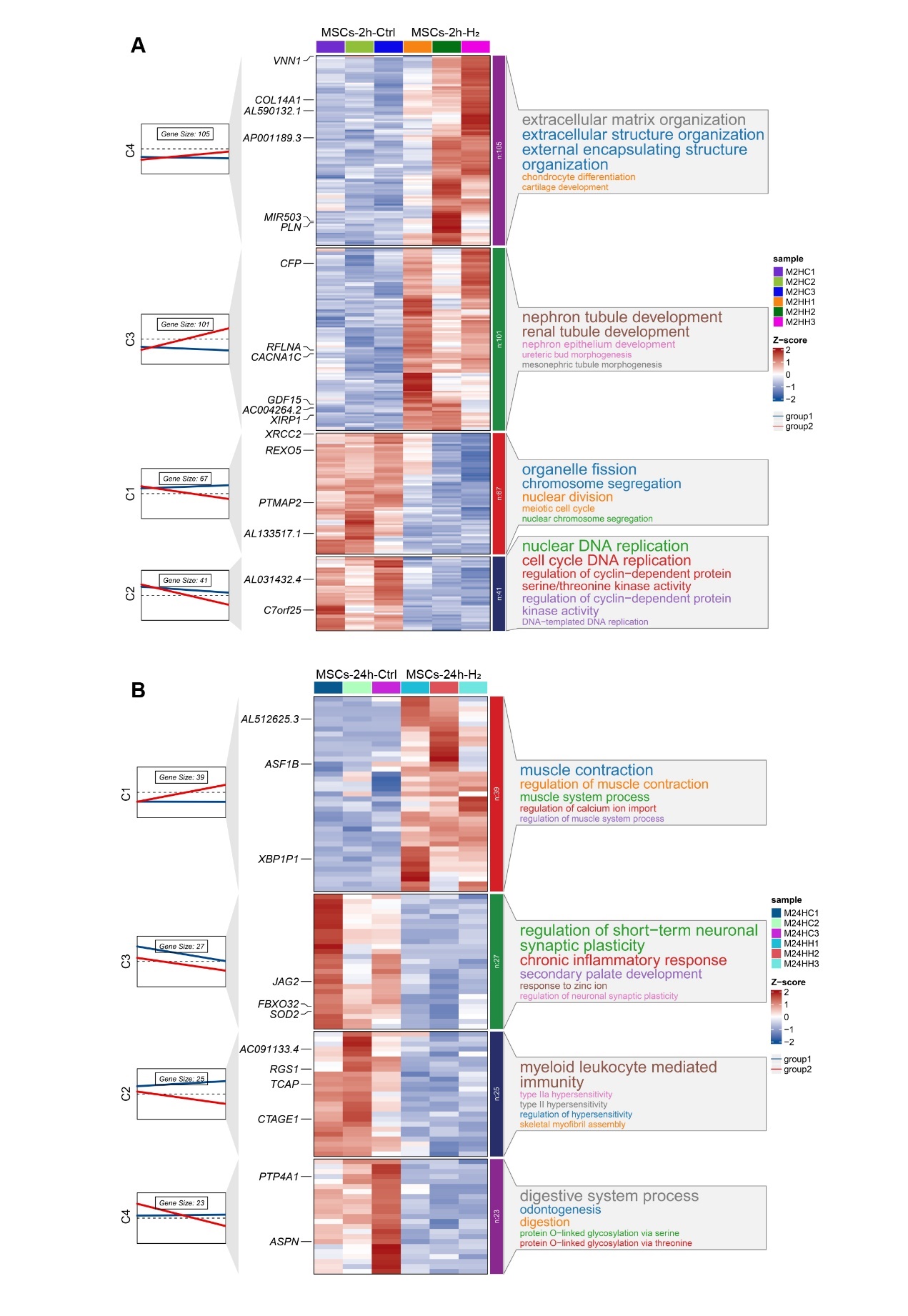
**Figure S7. Expression trends, heat maps, and significantly enriched Biological Process terms for each cluster of DEGs in MSCs after H_2_ treatment for 2 hours (A) and 24 hours (B).**

The left side shows a line plot of the expression trend of DEGs in each cluster of the control (blue line) and H_2_ group (red line); the left side of the rectangle shows the serial number of clusters, and the number of genes within each cluster (cluster size) is shown above the line plot; the center section presents heat maps of DEGs for each cluster, with color gradients from blue to red representing increasing expression levels; the right side shows the top 5 significantly enriched pathways from GO Biological Process enrichment analysis of DEGs in each cluster.

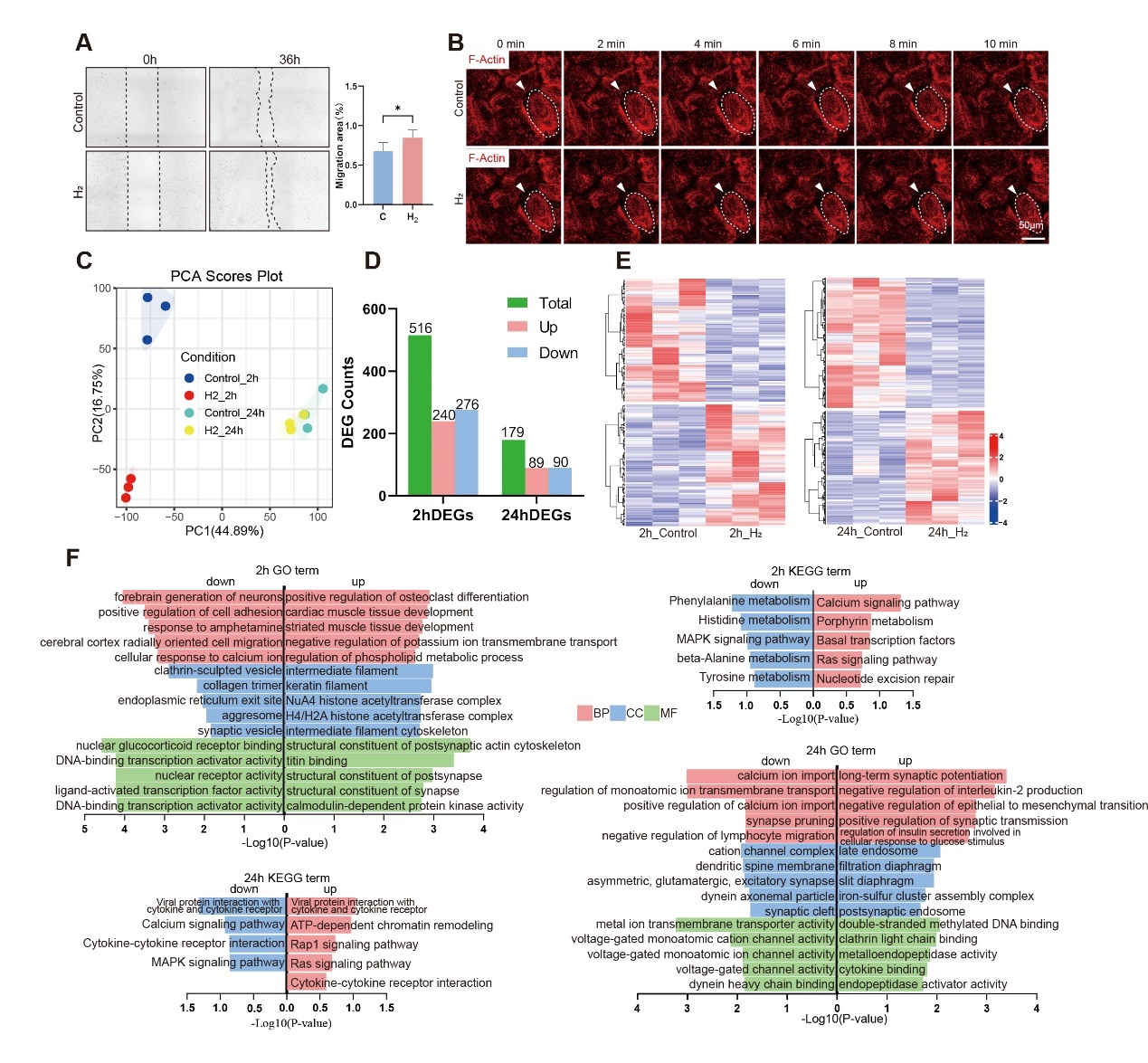
 **Figure S8. The impact of H_2_ on the migratory function of HUVEC cells and its effects on transcriptional expression.**

(A) Effect of H_2_ on the motion ability of HUVECs.
(B) The impact of H_2_ on the cytoskeletal structure of HUVECs.
(C) PCA diagram of sequencing data of each group.
(D) Differentially expression genes (DEGs) count results.
(E) Hierarchical clustering heat map of DEGs in HUVECs following 2-hour and 24-hour H_2_ treatment.
(F) H_2_-2h vs Control-2h and H_2_-24h vs Control-24h column charts for GO enrichment analysis and histogram of KEGG enrichment analysis. For B, using normal culture medium as the control group, changes in the cytoskeleton were captured over a 10-minute period. The medium was then immediately replaced with saturated hydrogen culture medium, ensuring consistent visibility, and changes were recorded for an additional 10 minutes. In the control group, there were no significant changes in the cytoskeleton during the initial 10 minutes. However, after switching to saturated hydrogen culture medium for 10 minutes, intercellular spacing increased, and the cytoskeleton underwent changes, showing a contraction trend. *p < 0.05

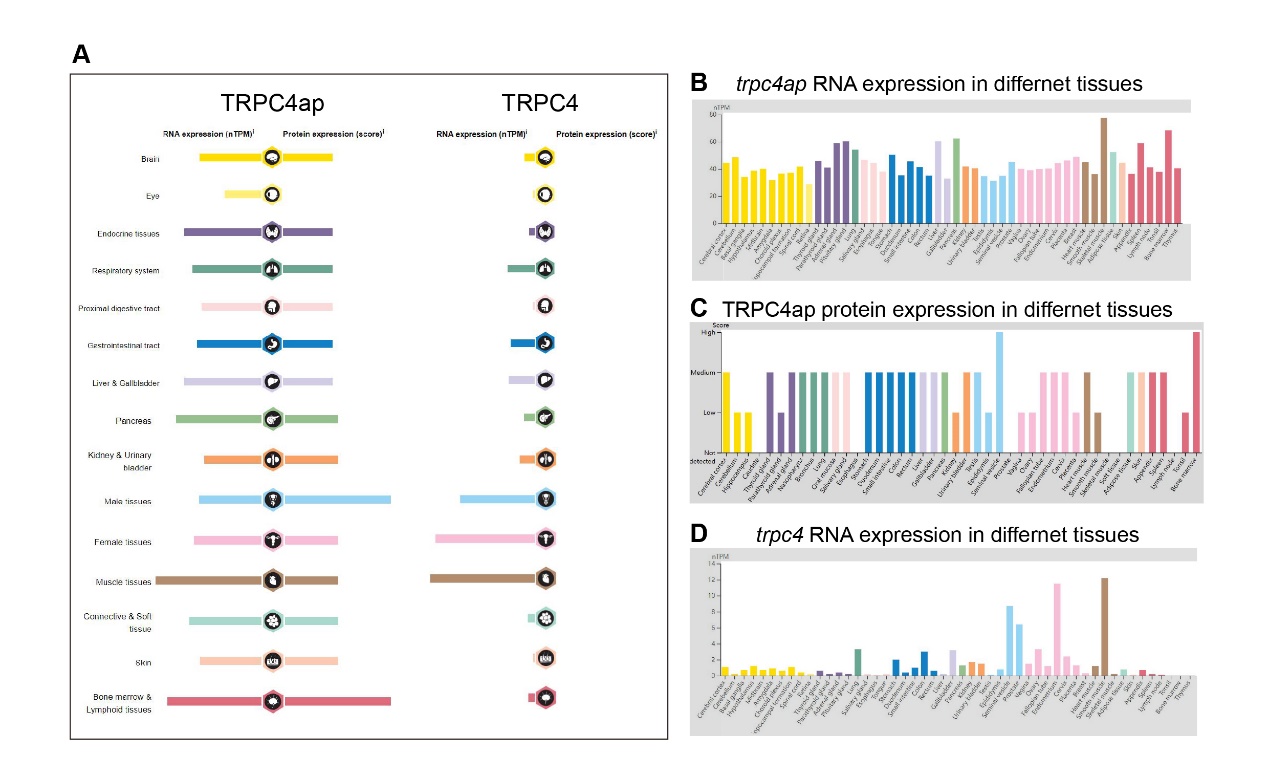

**Figure S9. Overview of TRPC4 and TRPC4ap expression based on data from the Human Protein Atlas.**

(A) depicts the expression levels of TRPC4ap and TRPC4 across various human organs, with RNA levels shown on the left and protein levels on the right.
(B) and (D) illustrate the RNA expression patterns of TRPC4ap and TRPC4 in distinct tissue types. C presents the protein expression distribution of TRPC4ap in multiple tissues. The images have been sourced, downloaded, and reorganized from the following URLs:

https://v19.proteinatlas.org/ENSG00000100991-TRPC4AP/tissue

https://v19.proteinatlas.org/ENSG00000133107-TRPC4/tissue

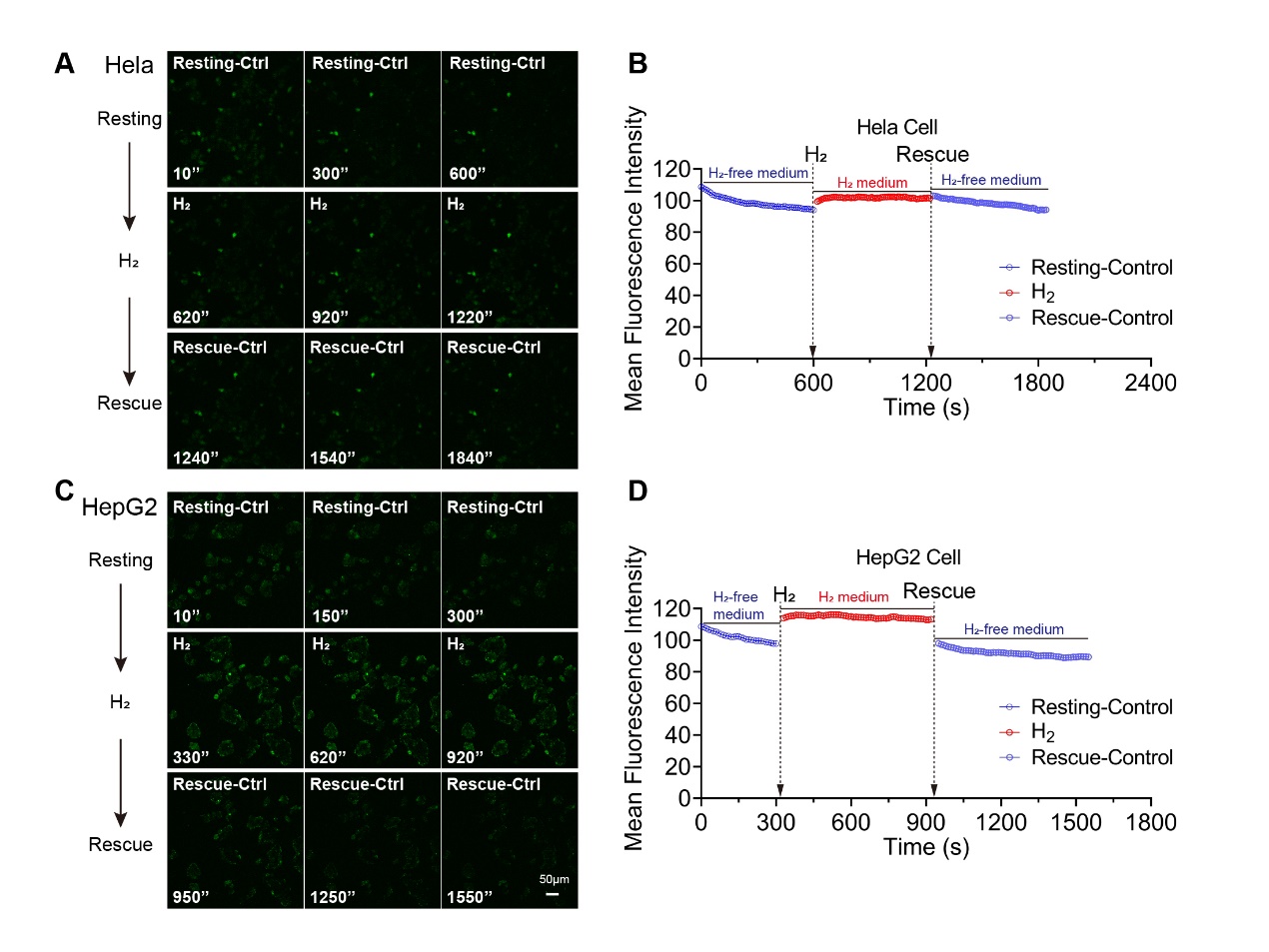

**Figure S10. The effect of H_2_-induced calcium influx is not evident in tumor cells Hela and HepG2.**

**(**A) & (C) Time-lapse images of the [Ca^2+^]i changes under H_2_-free, H_2_, and then H_2_-free conditions in Hela (A) and HepG2 (C) tumor cell lines. B & D Mean Fluorescence intensity changes in the three conditions of Hela (B) and HepG2 (D).
